# Interpretable Integration of CITE-seq RNA and ADT Profiles Without Explicit Modality-Weight Tuning via Tensor Decomposition-Based Unsupervised Feature Extraction

**DOI:** 10.64898/2026.04.18.719420

**Authors:** Turki Turki, Y-H Taguchi

**Affiliations:** Department of Computer Science, King Abdulaziz University, Jeddah 21589, Saudi Arabia; Department of Physics, Chuo University, 1-13-27 Kasuga, Bunkyo-ku, Tokyo 112-8551, Japan

**Keywords:** CITE-seq, T cells, enrichment analysis, unsupervised learning

## Abstract

CITE-seq jointly profiles cellular transcripts and surface proteins, but RNA and ADT modalities differ markedly in dimensionality, sparsity, and noise characteristics. We applied tensor-decomposition-based unsupervised feature extraction to paired CITE-seq data by constructing a gene × cell × protein tensor and performing HOSVD. The proposed workflow does not require explicit RNA/ADT modality-weight tuning or prior HVG-based gene filtering, and it provides cell-mode singular vectors together with post hoc unsupervised gene selection. Across ImmGen T-cell CITE-seq datasets, TD-derived cell representations preserved cell-type-related local structure and showed competitive kNN-based consistency compared with scMoMaT, a related factorization-based reference. However, ADT-only embeddings and Seurat WNN graphs often showed higher cell-type neighborhood consistency, indicating that TD-based UFE should not be interpreted as a replacement for marker-based or graph-based cell-type analysis. Enrichment analysis of TD-selected genes supported their biological plausibility but was interpreted as an exploratory check rather than proof of complete marker recovery. These results position TD-based UFE as a lightweight tensor-based unsupervised feature extraction framework for paired RNA/ADT data, rather than as a universally superior CITE-seq integration method.

## 1 Introduction

To completely understand the cellular processes, gene expression regulation, and complicated molecular mechanisms underlying RNA metabolism, not only single-omics layer measurements, but also multi-omics layer measurements must be performed at the single-cell level. During the last decade, single-cell RNA sequencing (scRNA-seq) has rapidly developed, enabling the mapping of cellular inhomogeneity in tissues, and has been employed as an analytical tool for investigating immunity against infectious diseases and cancers. However, single-cell transcriptomes do not always reflect protein expression on the cell surface or cell phenotypes. Delayed translation, post-transcriptional modification, alternative splicing, and differences in protein half-life and degradation rate can affect proteins on the cell surface, while maintaining the amount of mRNA. For example, in specific immune checkpoint inhibitor therapies, despite the increased ICOS protein on the cell surface, there are no changes in mRNA levels. This means that, in order to identify the uniqueness of cell types, differentiation trajectories, and signal transduction pathways, we need to bridge the gap between mRNA and protein.

CITE-seq (Cellular Indexing of Transcriptomes and Epitopes by sequencing) was developed to bridge this gap between transcript and surface-protein measurements. CITE-seq is a multimodal technology that simultaneously quantifies cell-surface protein expression using antibody-derived tags (ADTs) and intracellular mRNA expression in the same cells. In this procedure, antibodies conjugated with synthetic oligonucleotide barcodes are first bound to their target surface proteins. Cells with higher expression of a given surface protein bind more of the corresponding antibodies and consequently generate more ADT counts. The stained cells are then partitioned into droplets using a microfluidic platform such as the 10 × Genomics Chromium system, and sequencing libraries are prepared from both cellular mRNA and antibody-derived oligonucleotide tags.

The important advantage of CITE-seq is its ability to identify cell types because the number of surface protein molecules is much larger than that of intracellular mRNA and is hardly affected by dropout technology. In particular, surface proteins are convenient for identifying rare cell subpopulations that are difficult to identify using RNA measurements alone. It is also effective for detecting doublets (i.e., more than two cells are occasionally included in one droplet). However, the limitation of CITE-seq is that this technology can only detect surface proteins. If we attempt to barcode intracellular proteins, it also increases the possibility that the intracellular mRNA will start to leak, which decreases the amount of intracellular mRNA. If we can employ advanced methods such as DOGMA-seq, which enables the simultaneous measurement of mRNA, protein, and chromatin accessibility using digitonin (DIG) with streamlined processing, we can achieve more comprehensive analyses, although DOGMA-seq requires sophisticated technology.

To derive biologically meaningful insights from the massive multi-omics data measured by these advanced technologies, a highly advanced methodology is needed to integrate these datasets. Here, we summarize the statistical and computational difficulties encountered when attempting to integrate CITE-seq data, and outline the theoretical background, advantages, limitations, and benchmarks. The methods considered include graph-based, deep learning, matrix factorization, and mosaic integration approaches.

The most difficult aspect of the integration of CITE-seq data is that the statistical properties of protein and gene expression are distinct (Table 1). scRNA-seq measurements result in a large amount of sparse output, whereas protein measurements are dense and small. Because of the small number of ADT probes, the depth of ADT sequencing is large compared to that of scRNA-seq, which includes a large number of genes. In addition, since ADTs are measured using distinct antibodies with different affinities, batch effects cannot be ignored.

**Table 1.** Summary of difficulties.

| Modality | scRNA-seq (RNA) | CITE-seq (ADT) |
| --- | --- | --- |
| Dimension | High (20,000 - 30,000 ) | Low (20 - 200) |
| Property of data matrix | Sparse, many missing values | Dense, low missing values |
| Statistical model | Negative binomial distribution, zero-enhanced binomial distribution | Mixture model, continuous distribution |
| Major technological noises and artifact | Dropout, imbalanced depth of sequencing | Non-specific antibody binding, IgG batch effect |

Various methods have been proposed to address these difficulties (Table 2). Seurat

**Table 2.**
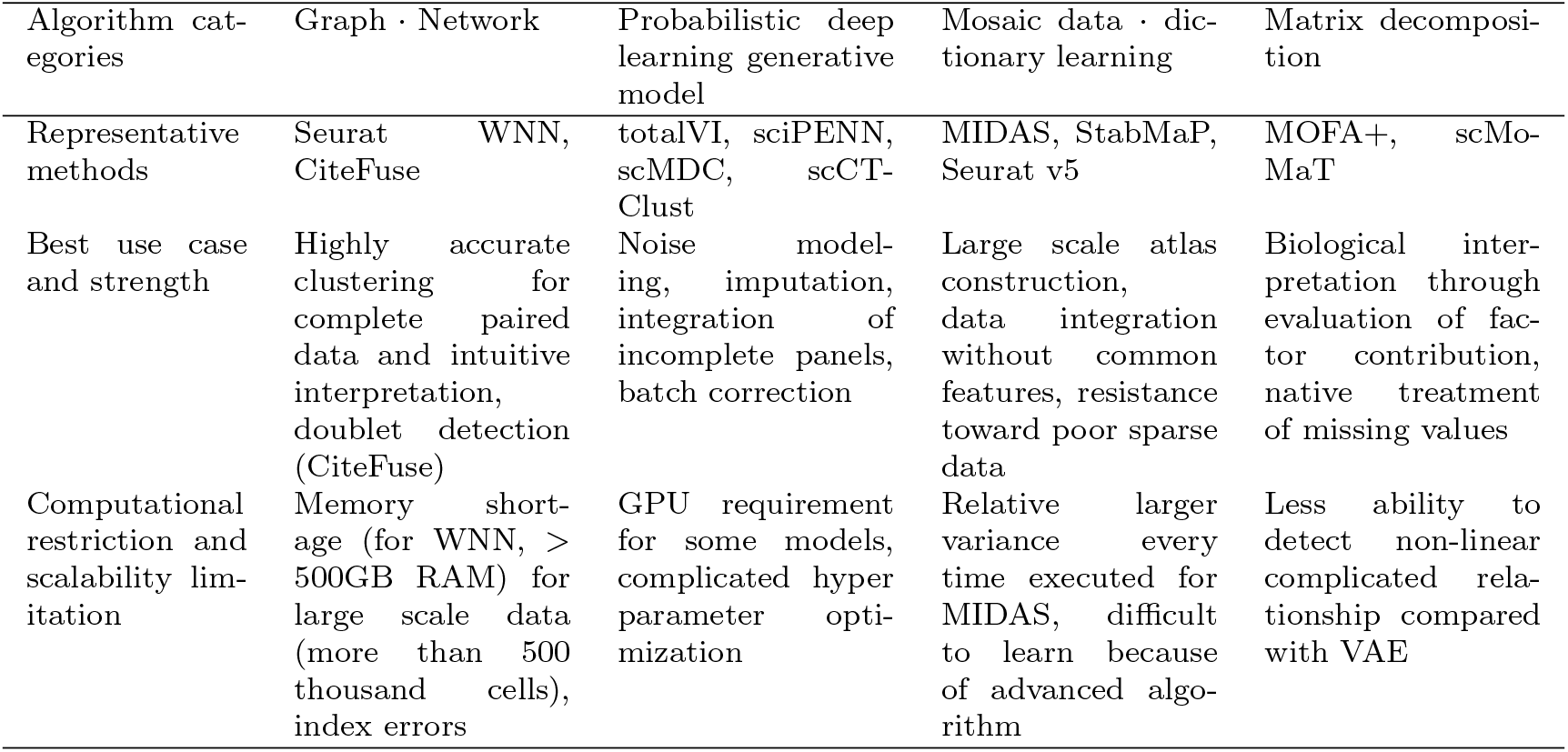
Summary of existing models.

WNN [1] dynamically evaluates the importance of RNA and protein modalities and addresses their modality weights. First, PCA is applied to the dataset, and a k-NN network is constructed. The variance of accuracy among neighbors when predicting cell types is evaluated, and the modality weight is tuned based on the variance (a higher variance results in a smaller modality weight). CiteFuse [2] constructs graph networks of RNA and proteins separately. The graph network of proteins is constructed based on similarity metrics computed using the propr package [3], whereas that of RNA is constructed based on the Pearson correlation coefficient. The two graphs are integrated using an iterative message-passing algorithm. Owing to the massive computational requirements for computing graph networks among cells, it is difficult to apply graph-based methods to too many cells (more than ≃ 10^5^ cells).

VAE (Variational Autoencoder)-based methods are designed to identify complicated non-linear relationships between cells. They aim to disentangle batch effects from the biological latent space and are also highly scalable. TotalVI [4] is a probabilistic latent model that represents gene and protein expressions as a combination of biological and technical factors. One remarkable point is the modeling of data uncertainty using Bayesian inference. TotalVI assumes that the noise property obeys the modality-specific parameters. This enables us to identify protein expression embedded in noisy signals. TotalVI is scalable to millions of cells. Single-cell imputation Prediction Embedding Neural Network (sciPENN) [5] is specific to the computational efficiency of the integration and inference of an incomplete dataset. sciPENN can deal with multiple profiles that do not always share the same protein species, and can predict protein expression using only gene expression. It is an order of magnitude faster than Seurat WNN and totalVI while maintaining the same accuracy. scCTClust [6] is yet another autoencoder-based technology, but deep-learning-based canonical correlation analysis is employed. It attempts to maximize the canonical correlation between proteins and genes. The fusion weights between proteins and genes are also optimized to obtain the best clustering consistency with cell type. Single-cell multi-omics deep clustering (scMDC) [7] is a multimodal deep learning model that targets clustering accuracy. It is composed of two independent autoencoders, one of which accepts gene and protein expression combined and the other accepts gene and protein expression separately. It simultaneously learns the latent space and generates clusters using a deep k-means algorithm. It can also correct batch effects based on a conditional autoencoder. When used with GPU, because of its linear scalability, it can handle massive large-scale multi-omics datasets.

Methods based on linear algebra have the advantage of biological interpretability, as it is clear which genes and proteins contribute to the outcomes. MOFA+ [8] decomposes the expression profiles into latent vectors and their weights. MOFA+ further decomposes the latent vector into parts common to genes and proteins and those unique to genes or proteins. MOFA+ also accepts missing data without pre-processing, which fills in the missing values. scMoMaT [9] can deal with a more complicated integration strategy by employing Matrix Tri-factorization, which enables the simultaneous embedding of genes, proteins, and cells.

However, for more extensive use cases (e.g., constructing an atlas database), there often appear cases where only genes or proteins are measured. The integration of these heterogeneous datasets is known as mosaic integration. More advanced methods must be developed for mosaic integration. MIDAS (Mosaic Integration and Knowledge Transfer of Single-cell Multimodal Data) [10] is a deep probabilistic framework designed for mosaic integration. The breakthrough in MIDAS lies in the alignment of modalities through self-supervised learning and the employment of information-theoretic latent disentanglement. StabMAP [11] employs mosaic data topology (MDT) and can make use of non-overlapping features, in contrast to traditional methods that can make use of only overlapping features. With multihop disjoint mosaic data integration, StabMAP can map cells without overlapping features in a common latent space. Seurat v5 [12] introduced bridge integration, which uses a molecular bridge to integrate multi-omics datasets, each of which has a unimodal distribution. This is based on the concept of dictionary learning where individual cells are regarded as elements in the dictionary. This enables Seurat v5 to integrate up to 33 million single cells.

In this study, we apply tensor decomposition (TD)-based unsupervised feature extraction (UFE) [13] to paired RNA/ADT CITE-seq data. TD-based UFE is related to factorization-based approaches such as scMoMaT in that it yields cell-, gene-, and protein-side representations. However, the present study does not aim to demonstrate universal superiority over existing CITE-seq integration methods. Rather, we examine whether a simple tensor-decomposition framework, without explicit RNA/ADT modality-weight tuning or prior HVG-based gene filtering, can provide a useful cell-side representation and post hoc unsupervised gene selection from the integrated RNA/ADT tensor.

The main contribution of this study is to demonstrate that the existing TD-based UFE framework can be extended to paired CITE-seq RNA/ADT data and can simultaneously provide a cell-side representation and post hoc unsupervised gene selection from the same integrated tensor, while clarifying its performance and limitations relative to scMoMaT, RNA-only, ADT-only, and Seurat WNN references.

## 2 Results

### 2.1 UMAP embedding using singular value vectors

After applying TD to *x*_*ijk*_, we obtained the singular value vector, 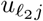, which is attributed to the *j*th single cell. UMAP was applied to the top ten 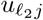, that is, 1 ≤ *ℓ*_2_ ≤ 10.

#### 2.1.1 GSE301271 (IGT36)

Figure 1 shows the various embeddings for GSE301271. The top left panel in Fig. 1 displays the results of TD (i.e., integration of gene and protein expression). The top right and bottom left panels show the UMAP embedding obtained from IGT36 for gene and protein expression, respectively (i.e., individual embedding for gene and protein expression). There are some notable points when UMAP embedding using TD (with integration) is compared to the two UMAP embeddings using either gene or protein expression (without integration). In the two UMAP embeddings using gene-only or protein-only expression, the same single-cell types were more scattered than those in the top left panel (using TD). This tendency was especially pronounced for CD4 cells; CD4 cells in TD embedding were clustered together, as shown in the top left panel of Fig. 1, but not in gene only- or protein only-based embeddings (top right and bottom left panels, respectively). The bottom right panel shows the results of totalVI taken from IGT36, which also integrates gene and protein expression. In the results of totalVI, CD4+ cells were more scattered than TD-based cells. In the UMAP visualization, the TD-derived representation showed visually coherent cell-type structure, particularly for major populations such as CD4 and CD8 cells. However, this visual impression was evaluated quantitatively in the subsequent section, because UMAP-based separation alone is not sufficient to establish superiority of the latent representation.

**Fig. 1.**
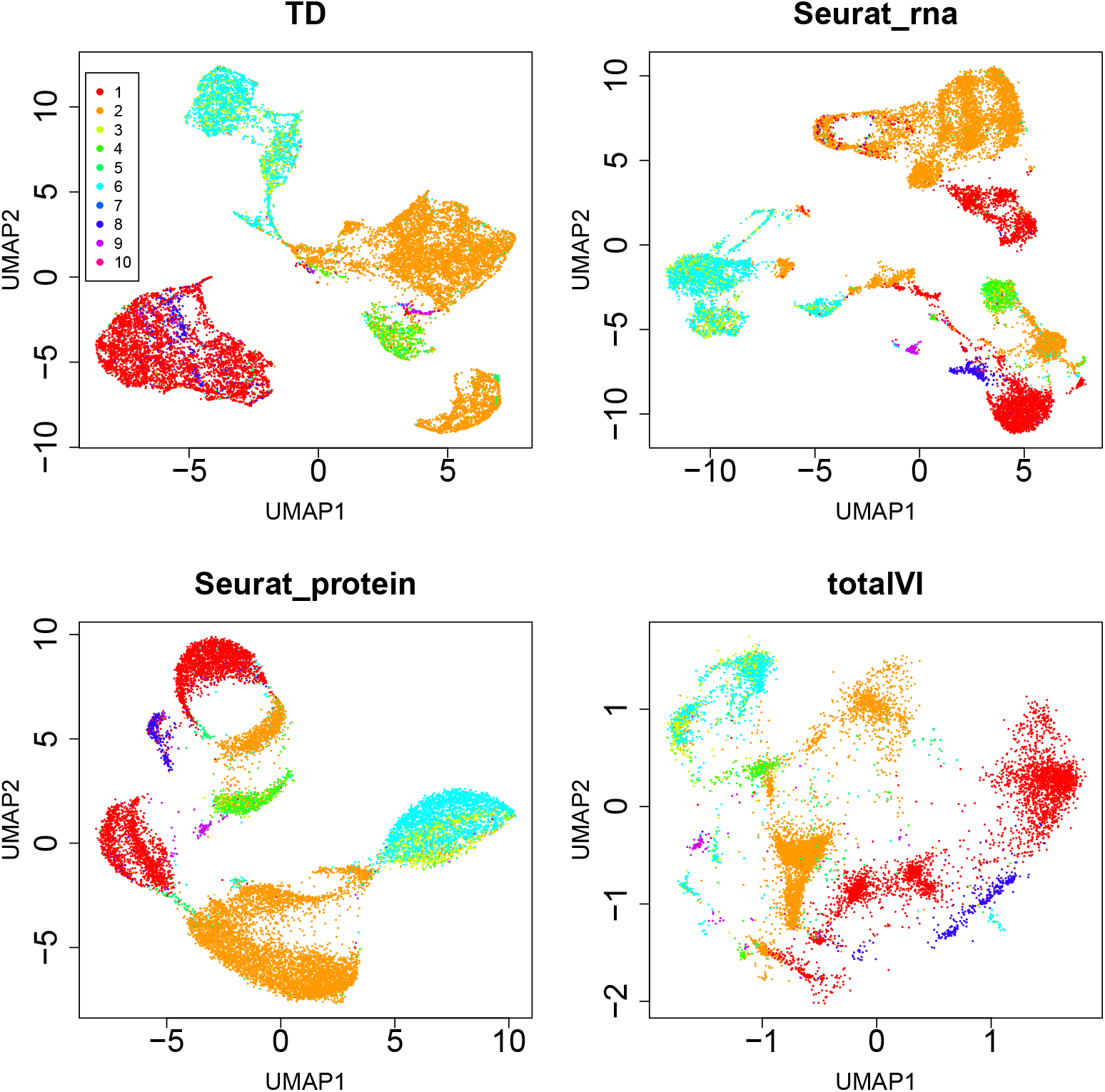
UMAP embedding for GSE301271 (IGT36). The coloring indicates level 1 classification of metadata (cell types). 1:CD4, 2:CD8, 3:CD8aa, 4:DN, 5:DP, 6:gdT, 7:nonT, 8:Treg, 9:Tz, 10:unclear.

We also assigned another classification labeling (organ) to these UMAP embeddings (Fig. 2). It is evident that integration did not improve the consistency between clusters in the UMAP embedding and labeling. This result is reasonable, since surface proteins, excluding tissue-specific markers, may not improve consistency with organ identity. Furthermore, the improved consistency between the UMAP embedding by TD and the annotated cell types in the top left panel of Fig. 1 is likely attributable to the integration of gene and protein expression, the latter of which is specifically designed to discriminate among cell types.

**Fig. 2.**
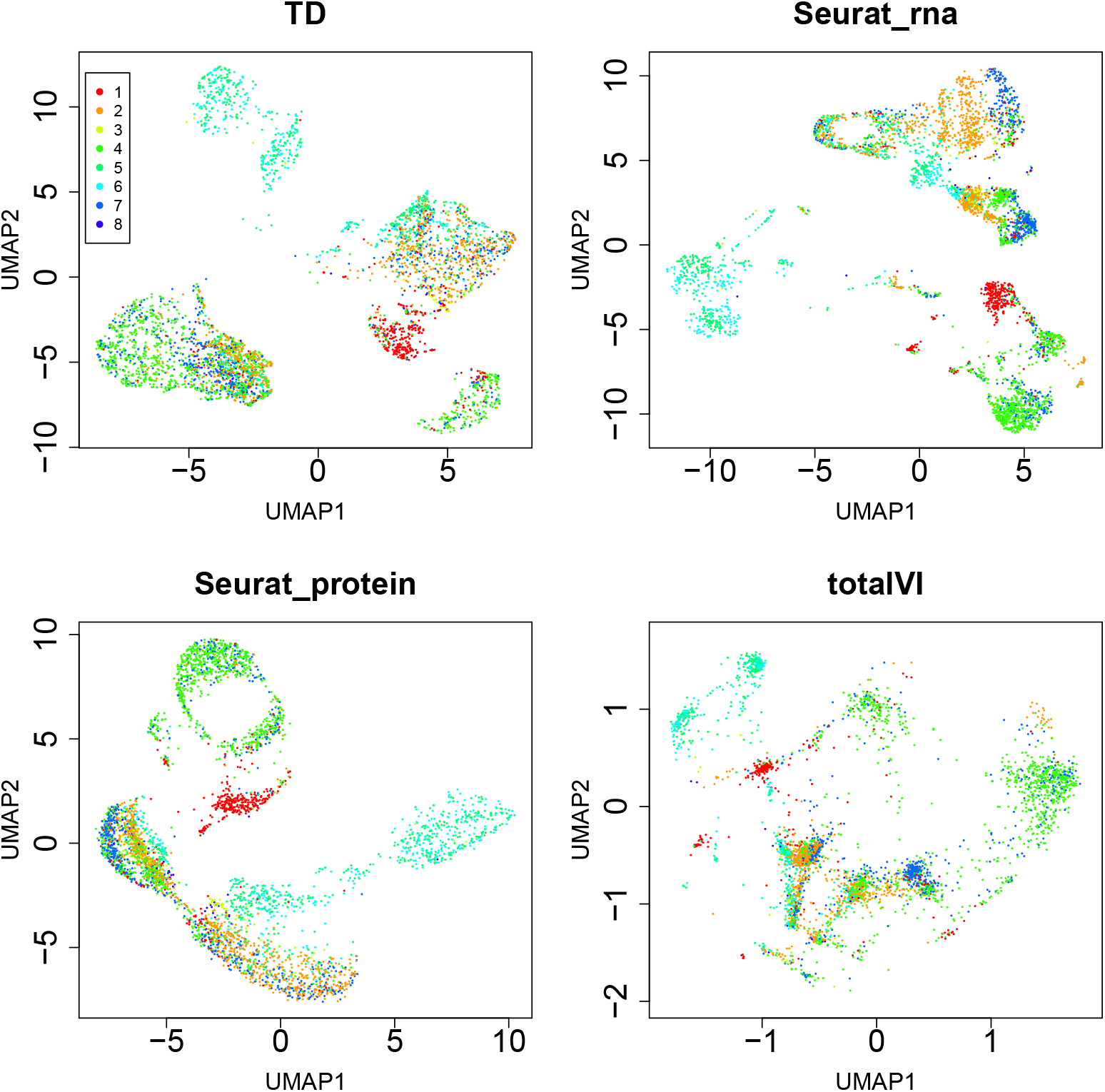
UMAP embedding for GSE301271 (IGT36). The coloring indicates organ classification of metadata (organ). 1:bone marrow, 2:lung, 3:prostate, 4:scLN, 5:SI IEL, 6:SI LP, 7:spleen, 8:sub-mandibular gland.

#### 2.1.2 GSE301960 (IGT38)

Figure 3 and 4 show the various embeddings of GSE301960 with cell type and organ specificity.

**Fig. 3.**
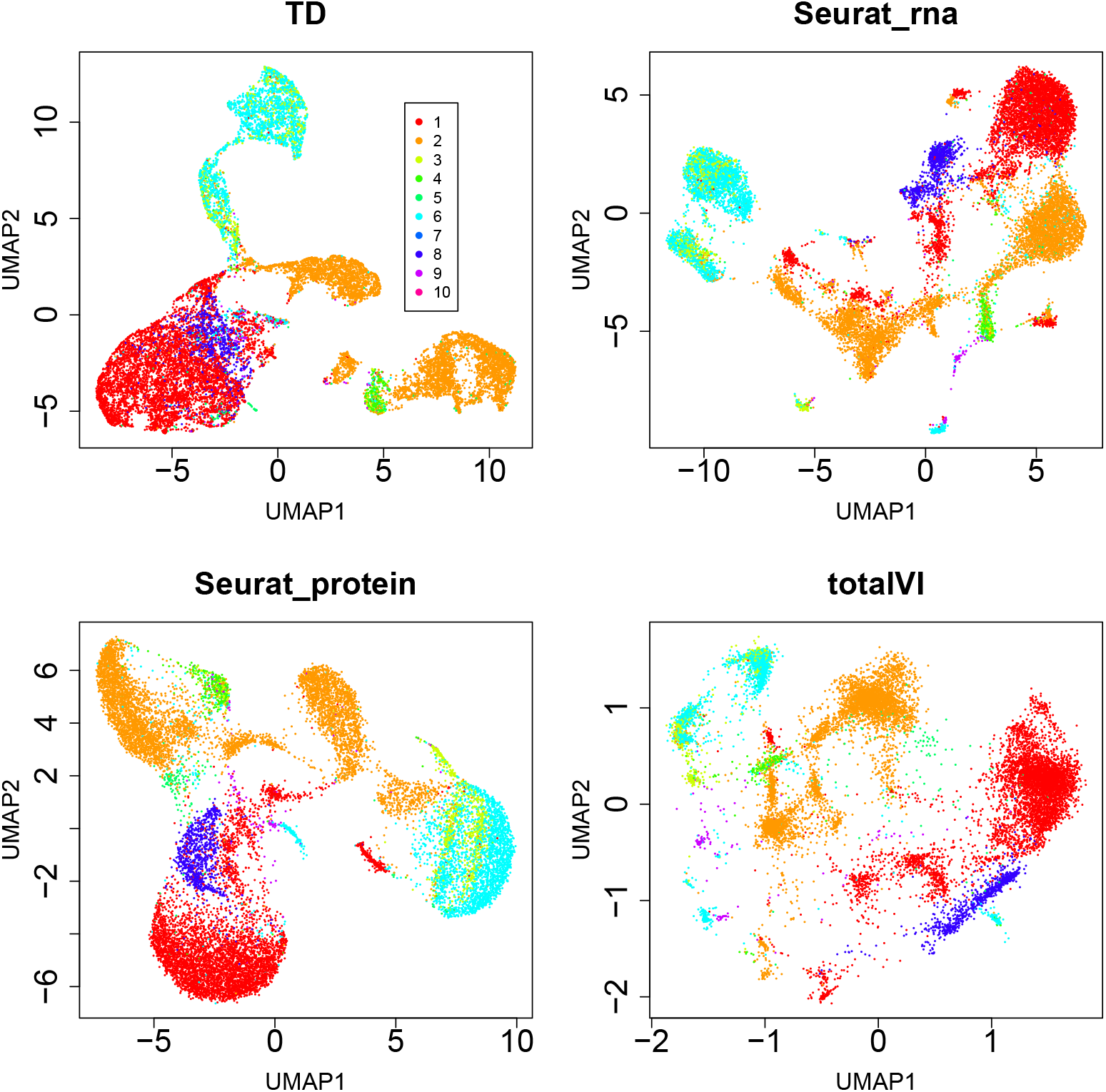
UMAP embedding for GSE301960 (IGT38). The coloring indicates level 1 classification of metadata (cell types) 1:CD4, 2:CD8, 3:CD8aa, 4:DN, 5:DP, 6:gdT, 7:nonT, 8:Treg, 9:Tz, 10:unclear.

**Fig. 4.**
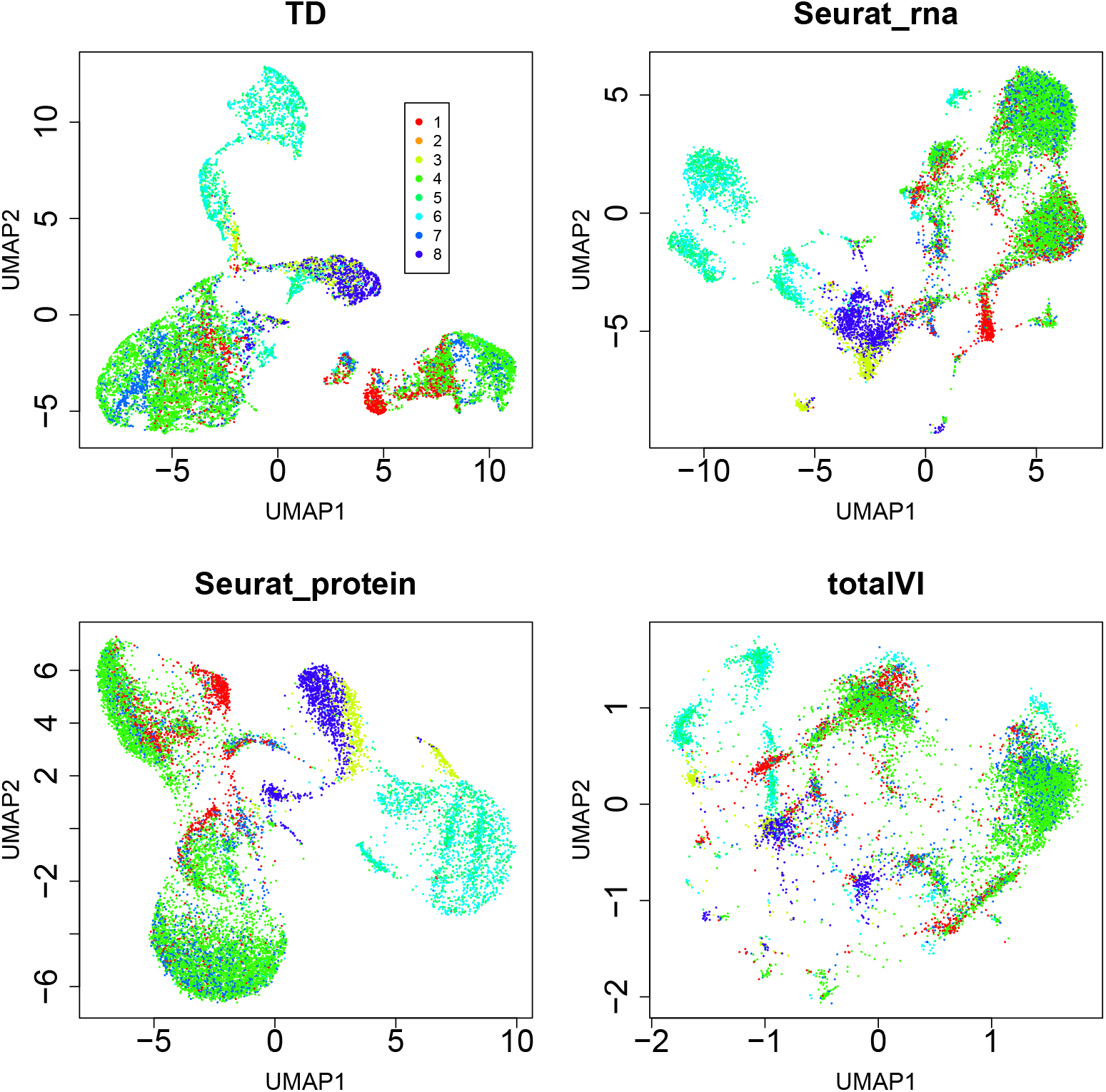
UMAP embedding for GSE301960 (IGT38). The coloring indicates organ classification of metadata (organ). 1:bone marrow, 2:lung, 3:prostate, 4:scLN, 5:SI IEL, 6:SI LP, 7:spleen, 8:sub-mandibular gland.

#### 2.1.3 GSE301961 (IGT40)

This dataset represents another additional time point under the same treatment conditions. Figure 5 and 6 show the various embeddings for GSE301961 with cell type and organ specificity for this dataset.

**Fig. 5.**
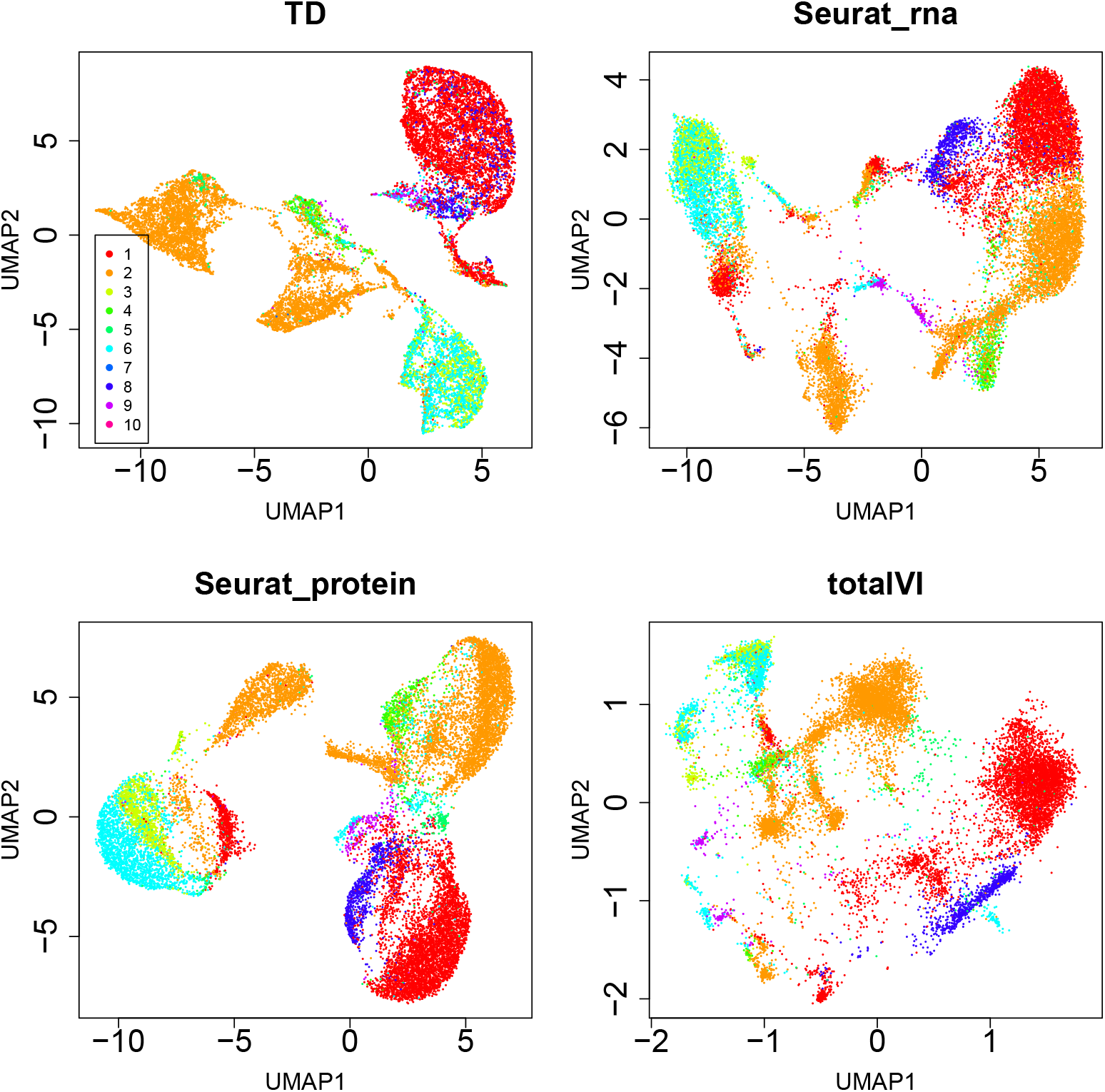
UMAP embedding for GSE301961 (IGT40). The coloring indicates level1 classification of metadata (cell types) 1:CD4, 2:CD8, 3:CD8aa, 4:DN, 5:DP, 6:gdT, 7:nonT, 8:Treg, 9:Tz, 10:unclear.

**Fig. 6.**
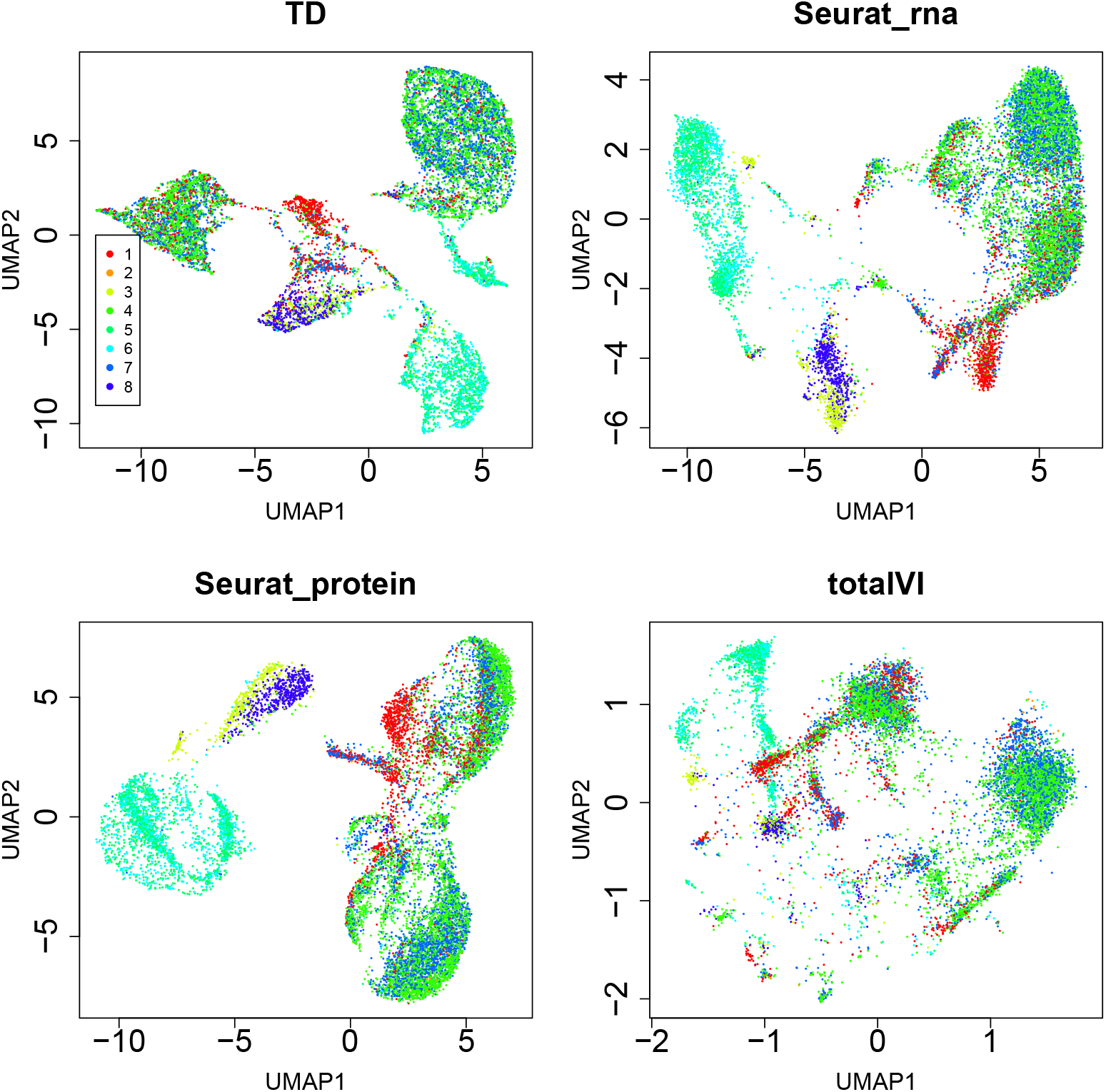
UMAP embedding for GSE301961 (IGT40). The coloring indicates organ classification of metadata (organ) 1:bone marrow, 2:lung, 3:prostate, 4:scLN, 5:SI IEL, 6:SI LP, 7:spleen, 8:sub-mandibular gland.

For GSE301960 and GSE301961, which represent additional time points under the same treatment condition, we observed qualitatively similar UMAP patterns. These visualizations suggested that the TD-derived UMAP structure was not limited to GSE301271. However, these observations were treated only as qualitative visualization results and were complemented by the quantitative kNN- and silhouette-based metrics described below.

#### 2.1.4 GSE281719 (IGT15)

We further examined GSE281719, a dataset obtained under a distinct experimental condition (Fig. 7). In this dataset, TD-derived UMAP visualization showed apparent separation of several major cell types. However, as shown in the quantitative comparisons, RNA-only and ADT-only references also performed strongly, and TD was not uniformly superior across metrics. Therefore, the UMAP results were interpreted as qualitative visualizations rather than as evidence of general superiority. Because this dataset was taken from only one organ, the submandibular gland, the comparison between UMAP embedding and organs is not shown, as it is not informative.

**Fig. 7.**
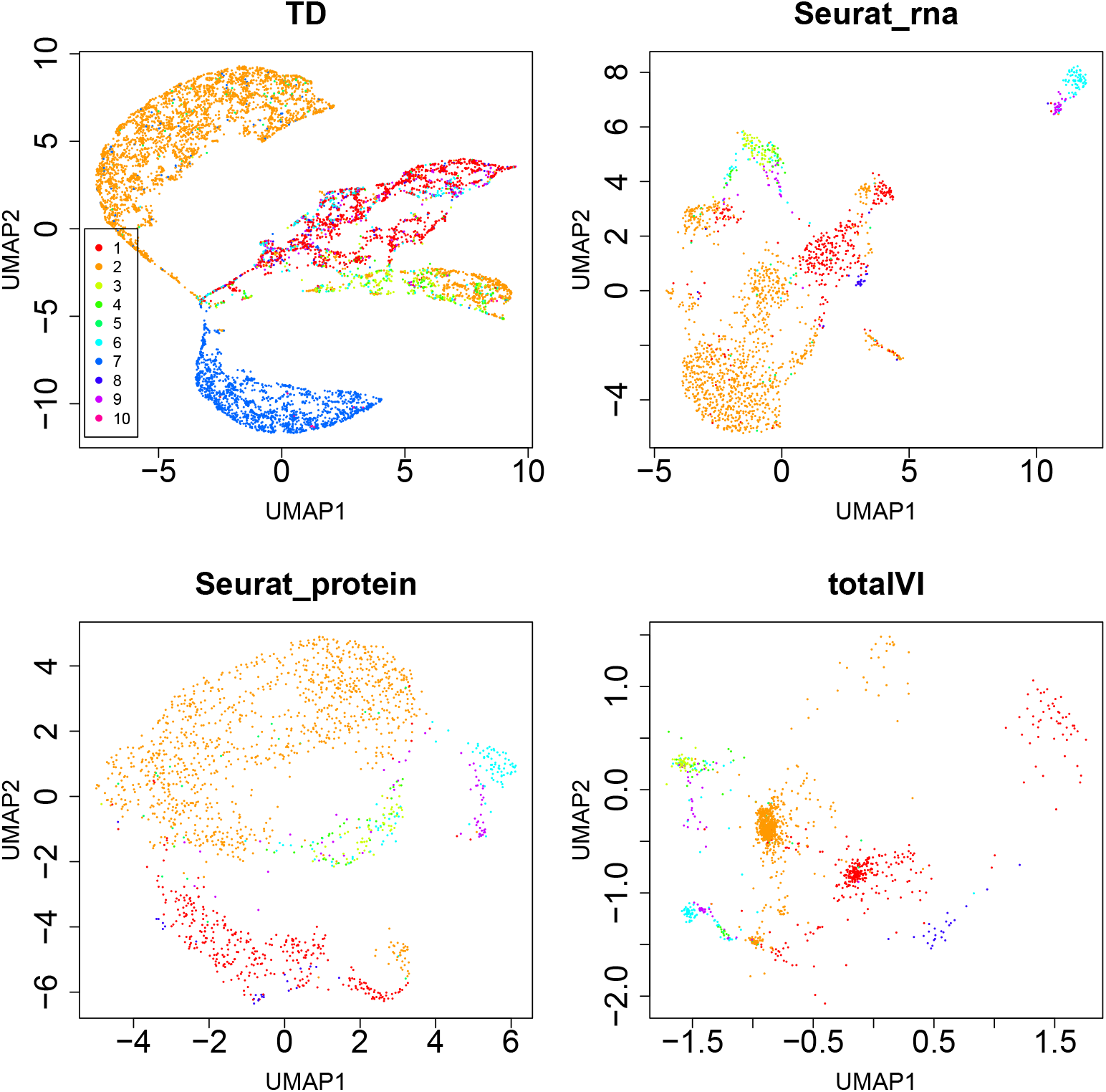
UMAP embedding for GSE281719 (IGT15). The coloring indicates level 1 classification of metadata (cell types). 1:CD4, 2:CD8, 3:CD8aa, 4:DN, 5:DP, 6:gdT, 7:nonT, 8:Treg, 9:Tz, 10:unclear.

#### 2.1.5 GSE283941 (IGT46)

TD-based UFE failed to produce a stable embedding for GSE283941, highlighting a limitation and a possible sensitivity to data quality or preprocessing conditions. Because we did not perform an independent quality-control analysis based on library size, detected genes, mitochondrial fraction, ADT background, doublet score, or annotation confidence, we do not interpret this failure as a formal QC criterion. Instead, we treat it as a limitation indicating that TD-derived embeddings may be sensitive to data quality or preprocessing conditions.

### 2.2 Quantitative comparison of cell-type neighborhood consistency

We next quantified the consistency between cell representations and annotated cell types using kNN purity, kNN majority-vote classification accuracy, macro-averaged metrics, and silhouette scores. For TD, RNA-only, ADT-only, and scMoMaT, these metrics were computed from Euclidean cell representations before UMAP. For Seurat WNN, graph-based kNN metrics were computed from the weighted nearest-neighbor graph, because WNN does not provide a conventional Euclidean latent embedding.

Therefore, WNN values were treated as graph-based reference metrics, and silhouette scores were not computed for WNN.

In the comparison with scMoMaT, TD showed higher overall kNN purity and kNN classification accuracy across the analyzed datasets, whereas macro-averaged and silhouette-based metrics varied by dataset and cell type (Table 3 and Table S1). In particular, Table 3 summarizes the dataset-level metrics, while Table S1 provides cell-type-specific classification accuracies. These results indicate that TD was competitive with scMoMaT as a related factorization-based reference, although TD did not uniformly outperform scMoMaT in all metrics.

**Table 3.** Various measures about the consistency between TD and scMoMaT embedding and cell types.

| Index | TD | scMoMaT | Interpretation |
| --- | --- | --- | --- |
| GSE301271 |  |  |  |
| kNN purity | 0.8396 | 0.7982 | TD > scMoMaT |
| macro kNN purity | 0.4462 | 0.4040 | TD > scMoMaT |
| kNN classification accuracy | 0.8876 | 0.8565 | TD > scMoMaT |
| macro kNN accuracy | 0.4778 | 0.4265 | TD > scMoMaT |
| silhouette | -0.1825 | 0.0272 | TD < scMoMaT |
| GSE301960 |  |  |  |
| kNN purity | 0.8230 | 0.7708 | TD > scMoMaT |
| macro kNN purity | 0.4493 | 0.3815 | TD > scMoMaT |
| kNN classification accuracy | 0.8743 | 0.8399 | TD > scMoMaT |
| macro kNN accuracy | 0.4866 | 0.3786 | TD > scMoMaT |
| silhouette | -0.1764 | -0.0134 | TD < scMoMaT |
| GSE301961 |  |  |  |
| kNN purity | 0.7900 | 0.7763 | TD > scMoMaT |
| macro kNN purity | 0.4500 | 0.4302 | TD > scMoMaT |
| kNN classification accuracy | 0.8350 | 0.8258 | TD > scMoMaT |
| macro kNN accuracy | 0.5034 | 0.4759 | TD > scMoMaT |
| silhouette | -0.1545 | 0.0913 | TD < scMoMaT |
| GSE281719 |  |  |  |
| kNN purity | 0.804 | 0.758 | TD > scMoMaT |
| kNN classification accuracy | 0.859 | 0.834 | TD > scMoMaT |
| macro kNN purity | 0.377 | 0.376 | TD $\simeq$ scMoMaT |
| macro kNN accuracy | 0.385 | 0.396 | TD < scMoMaT |
| silhouette | -0.015 | -0.032 | TD > scMoMaT |

We also compared TD with Seurat WNN, RNA-only, and ADT-only references. Seurat WNN showed higher graph-based cell-type neighborhood consistency than TD in most comparisons (Tables S2 and S3). RNA-only embeddings showed performance comparable to TD in some datasets and cell types, although the relative performance varied across datasets (Tables S4 and S5). ADT-only embeddings often showed higher cell-type consistency than TD, consistent with the fact that ADT panels contain surface markers directly related to annotated T-cell populations (Tables S6 and S7).

Together, these results indicate that TD-derived representations preserve cell-type-related local structure and are competitive with scMoMaT, but they do not support a claim that TD is generally superior to ADT-only, RNA-only, or WNN references.

### 2.3 Gene selection and enrichment analysis

In contrast to the standard procedure where gene selection must be performed prior to UMAP embedding, TD-based methods do not require any pre-selection of genes, and the obtained singular value vectors can be used for UMAP input as is (see Methods). Although this is a major methodological advantage of TD-based methods, we can also select genes even after embedding them to identify genes associated with the TD-derived RNA/ADT-integrated representation, namely the cell-type-related structure evaluated above. Using TD-based unsupervised FE (see Methods), we identified 6,707 genes for the GSE301271 dataset (a full list of the selected genes is available in the Supplementary Materials). Figure 8 shows the optimization process of 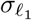 used for gene selection (for more details, see Methods).

**Fig. 8.**
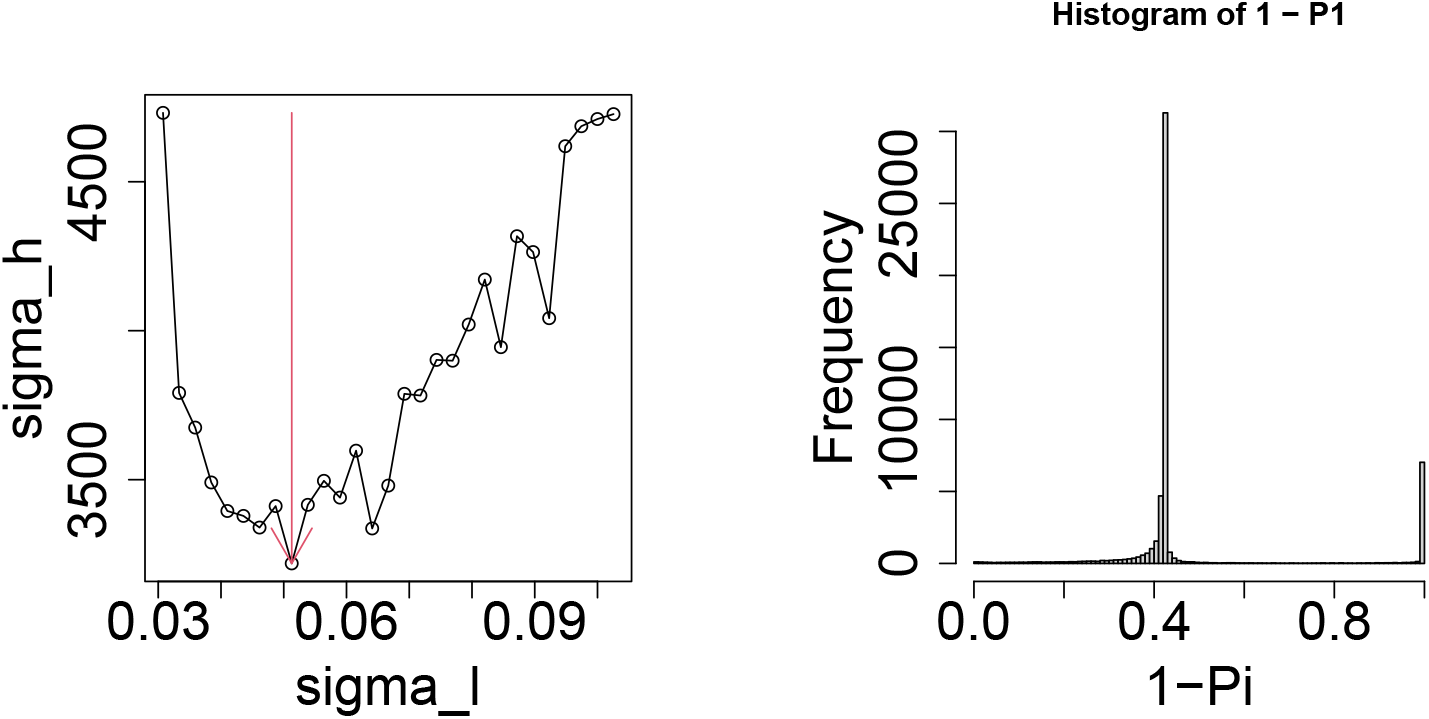
Optimization of 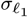 to select genes. Left: the dependence of *σ*_*h*_, which is the standard deviation of the histogram, upon 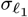. The vertical red arrow indicates the selected 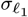. Right: Histogram of 1 − *P*_*i*_ using the optimized 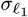. The right-end peak (i.e., *P*_*i*_ ∼ 0) corresponds to the bins including the selected genes.

To evaluate whether the TD-selected genes retained the TD-derived RNA/ADT-integrated representation, the entire procedure was repeated using only the 6,707 selected genes.

Figure 9 corresponds to Fig. 1. Although the three reference panels, except for the TD panel, are identical to those in Fig. 1, we retained them to facilitate comparison. Because the TD-derived UMAP based on the selected 6,707 genes was visually similar to that obtained using all genes, this result indicates that the TD-selected genes retained a substantial part of the RNA/ADT-integrated structure captured by TD-based UFE. It should also be noted that the gene selection was fully unsupervised, as no cell annotation information was used (for more details, see Methods).

**Fig. 9.**
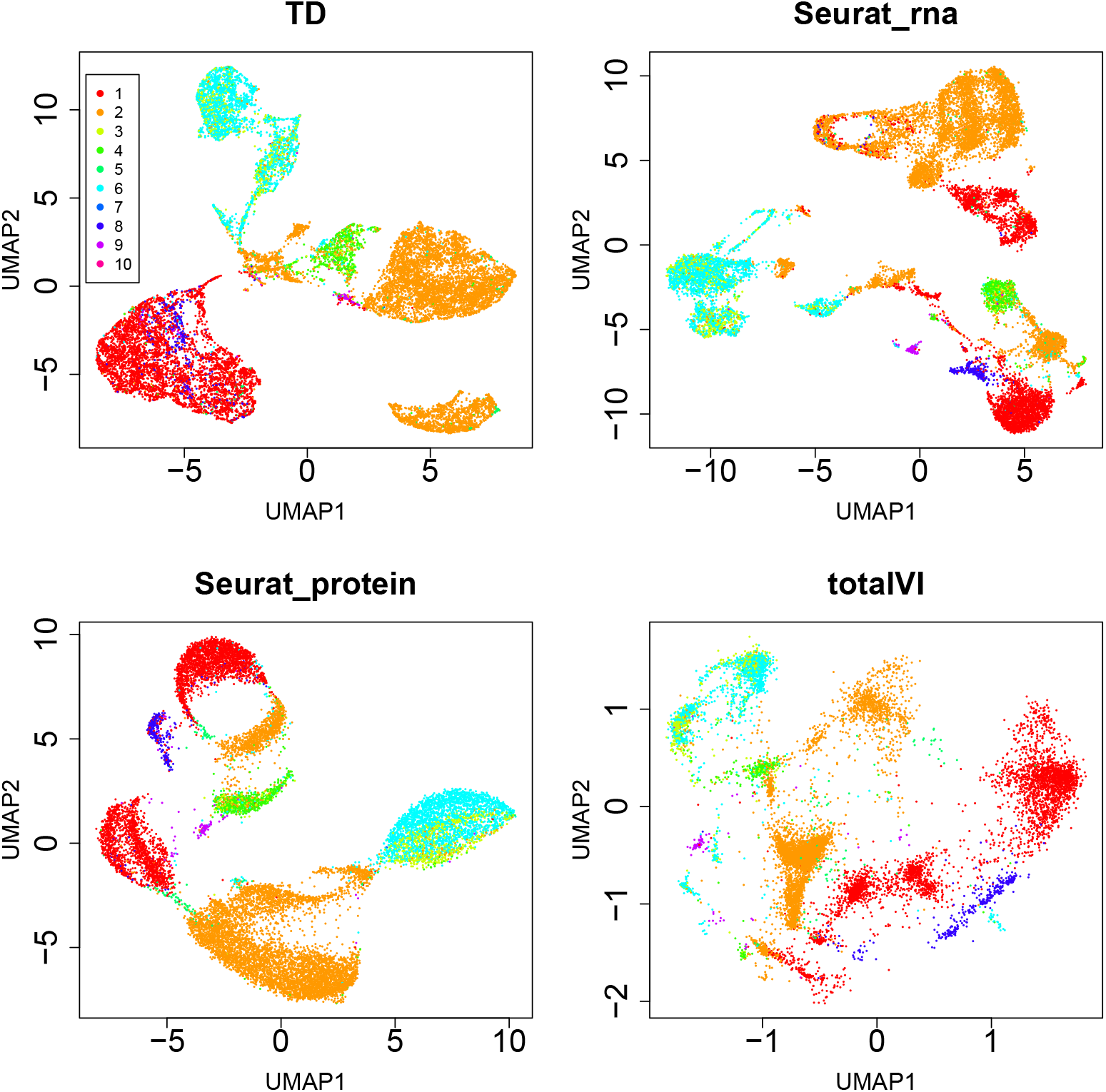
UMAP embedding for GSE301271 (IGT36) using only 6,707 genes selected by TD-based unsupervised FE. The coloring indicates level 1 classification of metadata (cell types). 1:CD4, 2:CD8, 3:CD8aa, 4:DN, 5:DP, 6:gdT, 7:nonT, 8:Treg, 9:Tz, 10:unclear.

Enrichment analysis was used as a biological plausibility check for the TD-selected genes. We did not interpret the selected genes as a complete set of canonical cell-type markers, nor did we use enrichment alone as evidence that TD-selected genes are superior to conventional feature-selection methods. Enrichment analysis was performed using Enrichr with its default background setting. Because a study-specific background gene universe corresponding to all genes used in TD was not explicitly specified, the enrichment results were interpreted as exploratory biological plausibility checks rather than as controlled over-representation tests. Some enriched terms were not directly related to T-cell biology. These terms may reflect broad reference signatures, shared marker genes, or nonspecific overlaps among large gene sets, and therefore were not interpreted as direct biological validation of TD-selected genes.

Table 4 lists the top 10 enriched terms for “Azimuth Cell Types 2021” category in Enrichr. These terms were consistent with the T-cell composition of the dataset. Table 5 lists the top 10 enriched terms for the “PanglaoDB Augmented 2021” category in Enrichr. These terms were also consistent with the T-cell composition of the dataset. Table 6 lists the top 10 enriched terms for another cell type enrichment category, “CellMarker Augmented 2021”, in Enrichr. Although several enriched terms were related to T-cell biology, some terms were not directly related to T cells. Therefore, these results were interpreted cautiously as exploratory enrichment patterns rather than direct validation of all selected genes. In addition to the cell type, we investigated tissue specificity, although the proteins were not designed to enhance tissue specificity. Table 7 lists the top enriched terms for the “Tabula Sapiens” category in Enrichr.

**Table 4.** Top 10 (either in TD or RNA-only. Since the union is presented, total number of terms can be larger than 10) enriched terms for “Azimuth Cell Types 2021” category in Enrichr for TD and RNA-only. If they do not match, they are shown as [TD,RNA-only]. Full list is available in the Supplementary Materials.

| Term | Overlap | P-value | Adjusted P-value |
| --- | --- | --- | --- |
| CD4+ Proliferating T CL0000896 | 13/13 | $6.73 \times 10^{-7}$ | $1.64 \times 10^{-4}$ |
| Proliferating Natural Killer CL0000623 | 13/14 | $6.49 \times 10^{-6}$ | $7.89 \times 10^{-4}$ |
| CD4+ Memory/Effector T CL0000624 | 10/10 | $1.79 \times 10^{-5}$ | $1.09 \times 10^{-3}$ |
| Cycling CL0008024 | 10/10 | $1.79 \times 10^{-5}$ | $1.09 \times 10^{-3}$ |
| CD8+ Naive T CL0000900 | [18,19]/24 | $[3.90, 5.98] \times 10^{-[5,6]}$ | $[1.90, 5.87] \times 10^{-[3,4]}$ |
| CD8+ Proliferating T CL0000906 | 10/11 | $1.37 \times 10^{-4}$ | $4.76 \times 10^{-3}$ |
| Proliferating Macrophage CL0000235 | 10/11 | $1.37 \times 10^{-4}$ | $4.76 \times 10^{-3}$ |
| Natural Killer T CL0000814 | [9,10]/11 | $[1.44, 1.37] \times 10^{-[3,4]}$ | $[2.33, 4.76] \times 10^{-[2,3]}$ |
| CD4 T CL0000624 | 9/10 | $3.73 \times 10^{-4}$ | $6.48 \times 10^{-3}$ |
| CD4+ Central Memory T 2 CL0000897 | 9/10 | $3.73 \times 10^{-4}$ | $6.48 \times 10^{-3}$ |
| CD4+ Effector Memory T 2 CL0001044 | 9/10 | $3.73 \times 10^{-4}$ | $6.48 \times 10^{-3}$ |

**Table 5.** Top 10 (either in TD or RNA-only. Since the union is presented, total number of terms can be larger than 10) enriched terms for “PanglaoDB Augmented 2021” category in Enrichr for TD and RNA-only. If they do not match, they are shown as [TD,RNA-only]. Full list is available in Supplementary material.

| Term | Overlap | P-value | Adjusted P-value |
| --- | --- | --- | --- |
| T Cells | [132,133]/165 | $[1.07, 1.30] \times 10^{-[34,35]}$ | $[1.89, 2.31] \times 10^{-[32,33]}$ |
| T Memory Cells | [112,113]/133 | $[1.31, 1.21] \times 10^{-[33,34]}$ | $[1.16, 1.07] \times 10^{-[31,32]}$ |
| Natural Killer T Cells | [99,101]/118 | $[1.38, 1.14] \times 10^{-[29,31]}$ | $[8.15, 6.74] \times 10^{-[28,30]}$ |
| Thymocytes | [98,99]/118 | $1.38 \times 10^{-[28,29]}$ | $[6.11, 4.89] \times 10^{-[27,28]}$ |
| NK Cells | [119,123]/157 | $[1.81, 8.72] \times 10^{-[27,31]}$ | $[6.39, 3.86] \times 10^{-[26,29]}$ |
| T Follicular Helper Cells | 92/110 | $2.23 \times 10^{-27}$ | $6.58 \times 10^{-26}$ |
| T Regulatory Cells | [95,96]/117 | $[2.85, 3.23] \times 10^{-[26,27]}$ | $[7.20, 8.16] \times 10^{-[25,26]}$ |
| T Cells Naive | 86/102 | $3.50 \times 10^{-26}$ | $7.73 \times 10^{-25}$ |
| T Helper Cells | [98,101]/148 | $[3.90, 5.53] \times 10^{-[16,18]}$ | $[7.66, 1.09] \times 10^{-[15,16]}$ |
| T Cytotoxic Cells | [70,68]/102 | $[4.37, 7.71] \times 10^{-[13,12]}$ | $[7.73, 1.05] \times 10^{-[12,10]}$ |
| Plasmacytoid Dendritic Cells | [86,93]/150 | $[1.75, 8.17] \times 10^{-[9,13]}$ | $[2.21, 1.45] \times 10^{-[8,11]}$ |

**Table 6.** Top 10 enriched terms for the “CellMarker Augmented 2021” category in Enrichr for TD and RNA-only. If they do not match, they are shown as [TD,RNA-only]. Full list is available in the Supplementary Materials.

| Term | Overlap | P-value | Adjusted P-value |
| --- | --- | --- | --- |
| Natural Killer T (NKT) cell:Fetal Kidney | [3237,2987]/4543 | 0.00 | 0.00 |
| Neural Stem cell:Hippocampus | [98,97]/99 | $[1.30, 1.29] \times 10^{-[45,43]}$ | $[6.40, 4.60] \times 10^{-[43,41]}$ |
| Regulatory T (Treg) cell:Liver | [280,292]/417 | $[1.81, 2.75] \times 10^{-[45,53]}$ | $[6.40, 1.48] \times 10^{-[43,50]}$ |
| Monocyte:Fetal Kidney | [451,448]/797 | $[2.95, 5.69] \times 10^{-[42,41]}$ | $[7.82, 1.22] \times 10^{-[40,38]}$ |
| Exhausted CD8+ T cell:Liver | [191,193]/260 | $[5.83, 1.78] \times 10^{-[40,41]}$ | $[1.24, 4.77] \times 10^{-[37,39]}$ |
| Vascular Stem cell:Blood | [95,96]/100 | $[5.48, 1.41] \times 10^{-[39,40]}$ | $[8.31, 2.17] \times 10^{-[37,38]}$ |
| Vascular Progenitor cell:Blood | [95,96]/100 | $[5.48, 1.41] \times 10^{-[39,40]}$ | $[8.31, 2.17] \times 10^{-[37,38]}$ |
| Oxyntic Stem cell:Stomach | [94,95]/99 | $[1.57, 4.08] \times 10^{-[38,40]}$ | $[1.66, 4.38] \times 10^{-[36,38]}$ |
| Oxyntic Progenitor cell:Stomach | [94,95]/99 | $[1.57, 4.08] \times 10^{-[38,40]}$ | $[1.66, 4.38] \times 10^{-[36,38]}$ |
| Progenitor cell:Stomach | [94,95]/99 | $[1.57, 4.08] \times 10^{-[38,40]}$ | $[1.66, 4.38] \times 10^{-[36,38]}$ |

**Table 7.** Top 10 enriched terms for the “Tabula Sapiens” category in Enrichr for TD and RNA-only. If they do not match, they are shown as [TD,RNA-only]. Full list is available in the Supplementary Materials.

| Term | Overlap | P-value | Adjusted P-value |
| --- | --- | --- | --- |
| Thymus-CD4-positive Helper T Cell | [90,91]/100 | $[4.25, 2.31] \times 10^{-[32,33]}$ | $[9.96, 5.41] \times 10^{-[30,31]}$ |
| Thymus-thymocyte | [90,92]/100 | $[4.25, 1.12] \times 10^{-[32,34]}$ | $[9.96, 5.24] \times 10^{-[30,32]}$ |
| Lymph Node-CD4-positive, Alpha-Beta Memory T Cell | [89,90]/100 | $[7.02, 4.25] \times 10^{-[31,32]}$ | $[2.99, 2.21] \times 10^{-[29,30]}$ |
| Lymph Node-naive Thymus-Derived Cd4-Positive, Alpha-Beta T Cell | [89,90]/100 | $[7.02, 4.25] \times 10^{-[31,32]}$ | $[2.99, 2.21] \times 10^{-[29,30]}$ |
| Lymph Node-CD4-positive Alpha-Beta T Cell | [89,90]/100 | $[7.02, 4.25] \times 10^{-[31,32]}$ | $[2.99, 2.21] \times 10^{-[29,30]}$ |
| Lymph Node-T Cell | [89,90]/100 | $[7.02, 4.25] \times 10^{-[31,32]}$ | $[2.99, 2.21] \times 10^{-[29,30]}$ |
| Lymph Node-type I NK T Cell | 89/100 | $7.02 \times 10^{-31}$ | $2.99 \times 10^{-29}$ |
| Skin-naive B Cell | [89,90]/100 | $[7.02, 4.25] \times 10^{-[31,32]}$ | $[2.99, 2.21] \times 10^{-[29,30]}$ |
| Skin-NK Cell | 89/100 | $7.02 \times 10^{-31}$ | $2.99 \times 10^{-29}$ |
| Thymus-dn3 Thymocyte | [89,90]/100 | $[7.02, 4.25] \times 10^{-[31,32]}$ | $[2.99, 2.21] \times 10^{-[29,30]}$ |
| Thymus-t Follicular Helper Cell | [89,90]/100 | $[7.02, 4.25] \times 10^{-[31,32]}$ | $[2.99, 2.21] \times 10^{-[29,30]}$ |
| Muscle-cd4-positive, Alpha-Beta T Cell | [88,89]/100 | $[1.05, 7.02] \times 10^{-[29,31]}$ | $[2.47, 2.35] \times 10^{-[28,29]}$ |

**Table 8.**
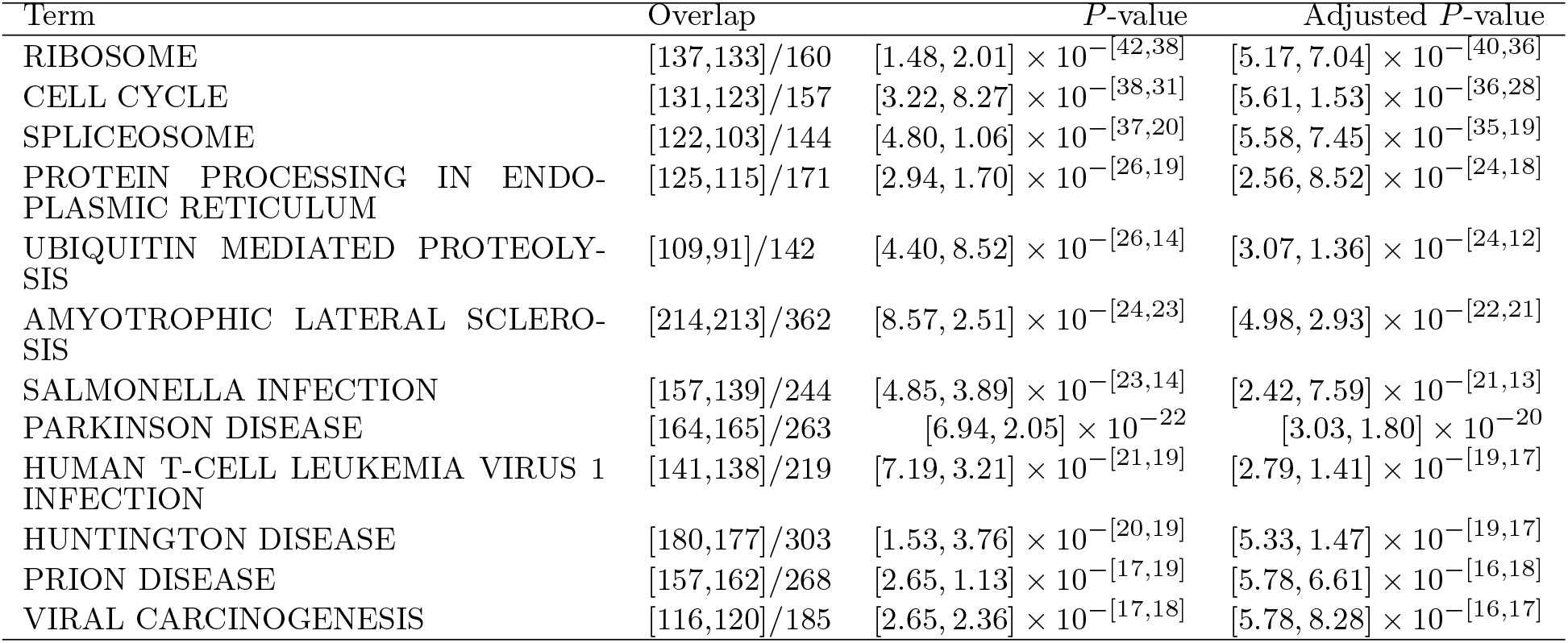
Top 10 (either in TD or RNA-only. Since the union is presented, total number of terms can be larger than 10) enriched terms for the “KEGG 2026” category in Enrichr for TD and RNA-only. If they do not match, they are shown as [TD,RNA-only]. Full list is available in the Supplementary Materials.

The enrichment of lymph-node-related terms was consistent with the tissue origin of GSE301271, which was measured in mediastinal lymph nodes.

These results support the biological plausibility of the selected genes, although they were interpreted only as exploratory enrichment patterns.

### 2.4 Diversity of latent space

Although we obtained the ten-dimensional space (i.e., the number of singular value vectors) used for UMAP embeddings consistent with cell-type-related structure, it remains unclear whether the latent space spanned by these ten dimensions is identical across each of the four experiments or shared among them. If they differ from each other, the latent space itself is experiment-dependent; otherwise, the latent space can represent the diversity of experiments. To observe this, we applied hierarchical clustering to 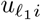 (for genes) or 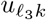 (for proteins), where there were 40 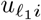 or 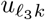 in total, respectively (Fig. 10). The results were distinct between genes and proteins. For genes, many 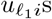 for GSE281719 (IGT15) did not cluster with any 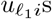 for the other experiments. This indicated that GSE281719 (IGT15) was distinct from the others. This was not surprising, because the other three were at distinct time points under the same treatment. For proteins, this type of isolation of GSE281719 (IGT15) from the other three isolates was not observed. This is because the number of proteins that aim to identify T cells is highly limited (180). This observation suggests that integrated analysis of genes and proteins in CITE-seq may lose the slight differences between distinct experiments, since variations that are not targeted by proteins may be overlooked. To provide a more quantitative assessment, we computed the mean square of some of the canonical correlations among the pairs of experiments:

**Fig. 10.**
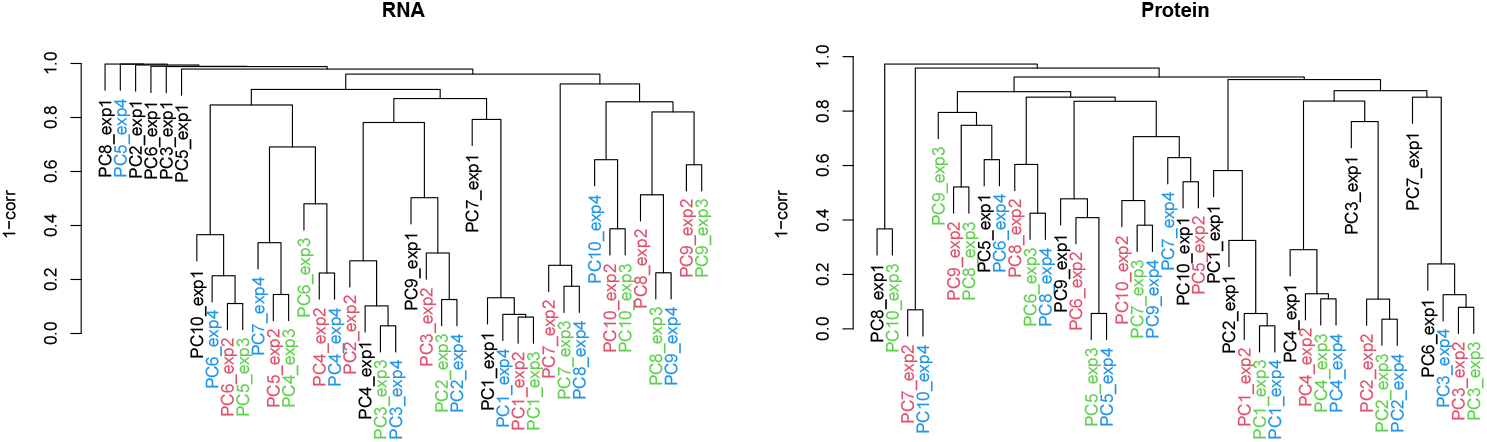
Hierarchical clustering of 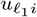 for genes (left) and 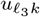 for proteins (right). Colors correspond to the four experiments. exp1(black):GSE281719 (IGT15), exp2(red):GSE301271 (IGT36), exp3(green):GSE301960 (IGT38) and exp4(blue):GSE301961 (IGT40).

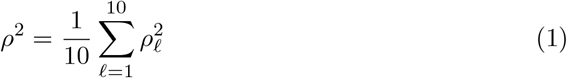

where *ρ*_*ℓ*_ is the *ℓ*th canonical correlation computed by the cancor function in R for 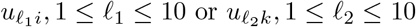 between a pair of experiments. *ρ*^2^ represents the ratio of the shared latent space between the two experiments (Table 9). As expected, GSE281719 (IGT15) had a smaller *ρ*^2^ value than the other three experiments for gene and protein expression. However, *ρ*^2^ among the three experiments excluding GSE281719 (IGT15) was nearly identical for proteins, but not for genes. The variation observed for genes is reasonable, because the time points of the three experiments were ordered as GSE301271 (IGT36) *<* GSE301960 (IGT38) *<* GSE301961 (IGT40). *ρ*^2^ between GSE301271 (IGT36) and GSE301960 (IGT38) or GSE301960 (IGT38) and GSE301961 (IGT40) should be larger than that between GSE301271 (IGT36) and GSE301961 (IGT40), which represents only gene expression. Protein expression clearly overlooked the differences in the time points among the three experiments. This suggests a limitation of the integrated analysis of gene and protein expression in CITE-seq experiments.

**Table 9.** Mean squared summation of the canonical correlation, *ρ*^2^. Upper half:genes; lower half: proteins

|  | GSE281719<br>(IGT15) | GSE301271<br>(IGT36) | GSE301960<br>(IGT38) | GSE301961<br>(IGT40) |
| --- | --- | --- | --- | --- |
| GSE281719 (IGT15) |  | 0.379 | 0.380 | 0.324 |
| GSE301271 (IGT36) | 0.436 |  | 0.801 | 0.673 |
| GSE301960 (IGT38) | 0.473 | 0.748 |  | 0.729 |
| GSE301961 (IGT40) | 0.433 | 0.737 | 0.716 |  |

## 3 Methods

All analyses were performed using the R statistical computing environment (version 4.3.1; https://www.r-project.org/) unless otherwise stated. TD-based UFE, HOSVD, gene selection, metric computation, and hierarchical clustering were performed using custom R scripts available at https://github.com/tagtag/TDbasedUFECITE-seq. Uniform Manifold Approximation and Projection (UMAP) was performed using the umap R package (version 0.2.10.0; https://CRAN.R-project.org/package=umap). RNA-only, ADT-only, HVG, and weighted nearest-neighbor analyses were performed using the Seurat R package (version 5.5.0; https://satijalab.org/seurat/). scMoMaT (single-cell Multi-omics Matrix Tri-factorization; version 0.2.2; https://github.com/PeterZZQ/scMoMaT) was run using Python version 3.12.3 (https://www.python.org/). Enrichment analysis was performed using the Enrichr gene-set enrichment analysis web server (accessed 24th July 2026; https://maayanlab.cloud/Enrichr/); the exact gene-set libraries used are specified below.

### 3.1 CITE-seq data set

We tested the following CITE-seq dataset using the GEO ID: GSE301271, GSE301960, GSE301961, GSE281719, and GSE283941 (Table 10). For each dataset, gene expression matrix, GSEXXXXXX matrix rna.mtx.gz, and protein expression matrix, GSEXXXXXX matrix protein.mtx.gz were downloaded and used for TD where GSEXXXXXXs are the corresponding GEO ID. They are a subseries of GEO ID GSE297097, from which various metadata sets are retrieved (see below), and also belong to the ImmGen T Open Source Project [14], from which metadata sets are retrieved (see below).

**Table 10.** Five CITE-seq data sets used in this study.

| GEO ID | Title | IGT ID | Number of |  |  |
| --- | --- | --- | --- | --- | --- |
|  |  |  | Genes | Proteins | Cells |
| GSE301271 | Pan-organ T cells at day 7 of acute LCMV Armstrong infection. | IGT36 |  |  | 17094 |
| GSE301960 | Pan-organ T cells at day 30 of LCMV Armstrong infection. | IGT38 |  |  | 17993 |
| GSE301961 | Pan-organ T cells at day 60 of LCMV Armstrong infection. | IGT40 | 55494 | 180 | 18222 |
| GSE281719 | Submandibular gland T cells in WT and PD1KO mice | IGT15 |  |  | 7937 |
| GSE283941 | Skin and lymph node CD8+ T cells after Vaccinia virus infection, DNFB | IGT46 |  |  | 7815 |

### 3.2 Meta data sets

Metadata sets were retrieved from various locations. For annotation of single cells (cell types and organs), GSE297097 annotation table 20260206 IGT1 104 cleaned.csv.gz was retrieved from GEO ID GSE297097. For cell ID, gene ID, and protein ID, GSE301271 cells.tsv.gz, GSEXXXXXX genes.tsv.gz, and GSEXXXXXX proteins.tsv.gz were retrieved from the corresponding GEO ID, GSEXXXXXX, such as GSE301271, GSE301960, GSE301961, GSE281719, or GSE283941. Finally, various UMAP embedding coordinates assigned to single cells included in the file IGTYY-cells-20260406.csv were retrieved from Rosetta2 on the ImmGen T Open Source Project website (https://rosetta.immgen.org/), where IGTYY denotes the corresponding IGT ID.

### 3.3 Generation of tensor

GSEXXXXXX matrix rna.mtx.gz and GSEXXXXXX matrix protein.mtx.gz were downloaded from the corresponding GEO ID, GSEXXXXXX. Because they are stored in a sparse matrix format, they were loaded into R using the readMM function in the matrix package [15].

We created a tensor *x*_*ijk*_ ∈ ℝ^*N ×M ×K*^ from the pairs of gene and protein expression matrices, *x*_*ij*_ ∈ ℝ^*N ×M*^ and *x*_*kj*_ ∈ ℝ^*K×M*^ where *i, j*, and *k* denote genes, cells, and proteins, respectively.

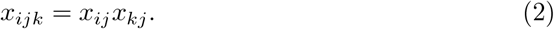

### 3.4 TD

TD was performed using HOSVD as follows:

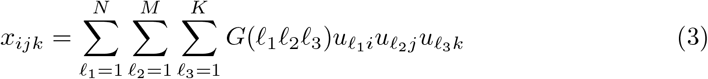

where *G* ∈ ℝ^*N ×M ×K*^ is the core tensor that represents the contribution of 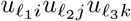 to 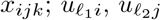, and 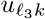 are the singular value matrices and orthogonal matrices. In this case, because the number of genes, *N* (∼ 5 × 10^4^), and cells, *M* (∼ 10^4^), are large, although *K*(∼ 10^2^) is modest, the number of nonzero elements in the generated tensor easily exceeds the maximum value representable by a 4-byte integer, even when a sparse matrix format is employed. Consequently, R is typically unable to manipulate a sparse matrix of this size. We therefore divided the large sparse matrix into a set of smaller sparse matrices, and TD was implemented using this set (the R code is available in the GitHub). See also the appendix.

### 3.5 UMAP

The UMAP was performed using the umap function in the umap package [16]. 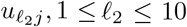 is employed as an input to UMAP. Other UMAP embedding for RNA-only, ADT-only and totalVI were retrieved from pre-computed data sets. See Meta data set subsection above and R code in GitHub.

### 3.6 Preprocessing and reference representations

For each dataset, RNA count matrices, ADT count matrices, and cell metadata were obtained from the corresponding GEO supplementary files. Cells were matched across RNA, ADT, and metadata tables using cell barcodes, and only cells present in all three tables were retained. The same matched cell set was used for TD, RNA-only, ADT-only, and scMoMaT comparisons whenever applicable. No cell-type annotation was used during tensor construction, HOSVD, or TD-based gene selection. No HVG-based gene filtering was performed before TD-based UFE.

For TD-based UFE and scMoMaT, the matched RNA and ADT count matrices obtained from the GEO supplementary files were used as input matrices without prior HVG-based gene selection. Zero entries were retained as zeros, and no imputation was performed before TD. The resulting RNA and ADT matrices were then used for tensor construction as described below.

For the quantitative RNA-only, ADT-only, and Seurat WNN reference analyses (i.e., metrics computation), we used the same Seurat object generated in the Seurat WNN workflow. In this workflow, the RNA assay was normalized using Seurat’s default log-normalization procedure, variable features were identified using Seurat’s standard variable-feature selection, the data were scaled, and PCA was performed. The RNA-only metrics were computed from the first 10 dimensions of the resulting RNA PCA reduction.

For the ADT assay, centered log-ratio (CLR) normalization was performed using Seurat’s ADT normalization workflow, followed by scaling and PCA. This ADT reduction was stored as APCA in the Seurat object. The ADT-only metrics were computed from the first 30 dimensions of the APCA reduction. Seurat WNN was then constructed from the RNA PCA and ADT APCA reductions using Seurat’s weighted nearest-neighbor procedure. Because Seurat WNN produces a weighted nearest-neighbor graph rather than a conventional Euclidean latent embedding, WNN was evaluated as a graph-based multimodal reference using graph-based kNN metrics.

Thus, RNA-only and ADT-only analyses were treated as single-modality reference representations derived within the same Seurat WNN preprocessing workflow, not as independent multimodal integration methods. When publicly available totalVI embeddings or UMAP coordinates from ImmGen/Rosetta files were used, they were treated as reference visualizations and not as fully controlled reimplementations under identical preprocessing.

### 3.7 Gene selections

#### 3.7.1 TD-based unsupervised FE

To determine the genes associated with the TD-derived RNA/ADT-integrated representation for deriving 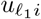 from *x*_*ijk*_, we employed TD-based unsupervised FE [13]. In TD-based unsupervised FE, *P* values are attributed to genes *i*, assuming that 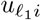 obeys a Gaussian (null hypothesis) as

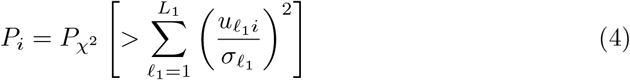

where 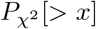 is the cumulative *χ*^2^ distribution; the argument is larger than *x* and 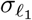 is the standard deviation to be optimized as follows: Gene selection was performed by selecting genes associated with *P* -values adjusted by the BH criterion [13] to be less than 0.01.

To optimize 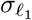, we attempted to make the histogram of 1 − *P*_*i*_ after excluding the selected indices *i*s as flat as possible, because the histogram of 1−*P*_*i*_ should be uniform when the null hypothesis is true. The procedure for optimizing 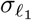 is as follows:

1. Set the same value *σ*_0_ to 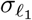.
2. Attribute *P*_*i*_ to the corresponding indices *i* using eq. (4) with setting *L*_1_ = 10.
3. Compute the histogram of 1 − *P*_*i*_ after excluding indices *i* associated with adjusted *P* -values less than 0.01.
4. Compute the standard deviation of histogram, *σ*_*h*_, as

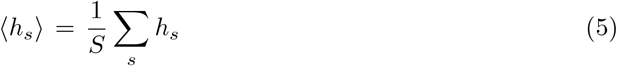

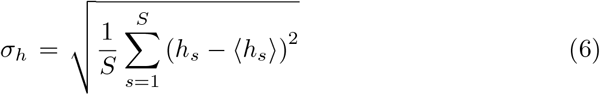

where *h*_*s*_ is the histogram of 1-*P*_*i*_ at the *s*th bin and *S* is the total number of bins (Usually, *S* = 100).
5. Update *σ*_0_ so as to reduce *σ*_*h*_.
6. If *σ*_*h*_ does not converge, go back to step 2.

Importantly, this process is fully unsupervised because we did not make use of any annotation information of the cells.

#### 3.7.2 Highly variable genes (HVG)

HVGs were selected using RNA counts provided by Seurat. The R scripts used for this analysis are available in the GitHub repository.

#### 3.7.3 RNA only

RNA-only-selected genes were selected using the feature loadings of the RNA PCA reduction generated in the Seurat object used for RNA-only metric computation. Specifically, the R script ranked genes according to their contributions to the RNA PCA space used for the RNA-only reference metrics, and the same number of genes as the TD-selected gene set was retained. This RNA-only-selected gene set was used as a reference for comparison with TD-selected genes. The R scripts used for this analysis are available in the GitHub repository. The precomputed RNA-only UMAP coordinates shown in the figures were used only as reference visualizations and were not used for RNA-only gene selection.

### 3.8 Enrichment analysis

Enrichment analysis was performed using Enrichr [17] with its default background setting. Because a study-specific background gene universe was not explicitly specified, the enrichment results were interpreted as exploratory biological plausibility checks rather than controlled over-representation tests. KEGG [18] pathway enrichment was evaluated using the KEGG 2021 Human gene-set library provided through Enrichr.

### 3.9 Hierarchical clustering of singular value vectors

Hierarchical clustering of 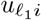 and 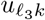 was performed using `hclust` function in R with “method=‘ave’ “ option. The pairwise distance was 1 minus the absolute value of the Pearson correlation coefficient.

### 3.10 scMoMaT

scMoMaT was applied to GSEXXXXXX matrix rna.mtx.gz and GSEXXXXXX matrix protein.mtx.gz using the default settings (4,000 iteration steps (epochs)). The R code used for this analysis is available on GitHub.

### 3.11 Seurat WNN

Seurat WNN was applied to GSEXXXXXX matrix rna.mtx.gz and GSEXXXXXX matrix protein.mtx.gz using the Seurat WNN workflow described above. The WNN graph was constructed from the RNA PCA and ADT APCA reductions, and graph-based kNN metrics were computed from the resulting weighted nearest-neighbor graph. The R code used for this analysis is available on GitHub.

### 3.12 Metric computation

All metrics were computed using R scripts available in the GitHub repository.

## 4 Discussion

There are several advantages of TD-based methods. First, TD-based methods did not result in cell loss. In standard approaches, low-quality cells are usually discarded prior to analysis. TD can be applied without prior cell removal in the present workflow. Previously, we successfully demonstrated that TD-based unsupervised FE can integrate a single-cell multi-omics dataset full of missing values without filling in the missing values in the preprocessing [19]. Additionally, in TD-based approaches, no gene selection is required prior to UMAP because TD itself generates singular value vectors to be used as inputs for UMAP. Genes can also be selected after UMAP because TD is a linear method. In typical nonlinear methods, it is impossible to remove some parts of genes after obtaining the latent space. In TD, this is straightforward because TD is a linear method and is completely reversible.

Moreover, regarding the implementation of TD, in this study, TD for the product of two large sparse matrices can be implemented. Since the size of the tensor is *N* × *M* × *K*, when the sample size *M* is large, then *N* × *M* × *K* can also become large because the number of features *N* and *K* is usually large. In this case, the sample index is usually summed to reduce the size of the tensor to that of a matrix *N* × *K* [13]. Nevertheless, in this study, handling a large tensor of size up to *N* × *M* × *K* becomes feasible by splitting it into a set of smaller tensors. This enables the application of TD-based unsupervised FE to a wide range of single-cell experiments where the sample size can be large.

As mentioned in the Introduction, the advantage of TD-based unsupervised FE over scMoMaT is that TD-based unsupervised FE does not require any tuning parameters that scMoMaT has. The objective function to be minimized in scMoMaT is:

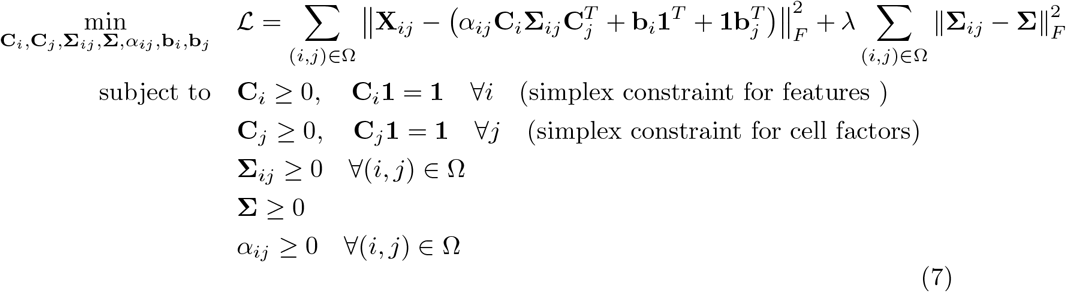

where *X*_*ij*_ is the expression of the *i*th feature (i.e., gene or protein) in the *j*th cell, Σ_*ij*_ is a matrix relating *i* to *j, C*_*i*_ and *C*_*j*_ are latent vectors assigned to features (i.e., genes or proteins) and cells, respectively; ***b***_*i*_ and ***b***_*j*_ are bias vectors; Ω represents a complete set of indices attributed to cells, genes, and proteins.

TD-based UFE does not have tuning parameters corresponding to *λ, α*_*ij*_, *b*_*i*_, or *b*_*j*_ in the scMoMaT formulation. Among these parameters, the most critical difference between scMoMaT and TD-based UFE is the presence of *α*_*ij*_, which controls the relative weight between gene and protein measurements in scMoMaT. TD-based UFE does not have a term corresponding to *α*_*ij*_, because each tensor element *x*_*ijk*_ is defined as the product of two matrices, *x*_*ij*_*x*_*kj*_, so that each entry contains both gene and protein expression in the same multiplicative form. Thus, no additional explicit rebalancing parameter between genes and proteins is introduced in TD-based UFE, despite the large difference in the numbers of genes and proteins considered.

This formulation is consistent with previous applications of TD-based UFE to the integration of two omics profiles with very different feature numbers, such as genes (∼ 10^4^) and miRNAs (∼ 10^3^), or genes (∼ 10^4^) and methylation sites (∼ 10^6^) [13]. These previous results motivated the use of the same product-form tensor construction for CITE-seq RNA/ADT integration without introducing additional tuning parameters to rebalance proteins and genes. Because the choice of *α*_*ij*_ can affect the resulting representation, TD-based UFE avoids one source of explicit modality-balancing dependence that is present in some matrix-factorization approaches. Under the tested settings, TD-based UFE produced embeddings comparable to scMoMaT while offering a simpler workflow. However, this does not imply general superiority across all CITE-seq datasets.

Figure 11 shows the UMAP embeddings obtained using scMoMaT. The top-left, top-right, bottom-left, and bottom-right panels in Fig. 11 correspond to the TD-based UMAP panels for GSE301271, GSE301960, GSE301961, and GSE281719 shown in Figs. 1, 3, 5, and 7, respectively. These visualizations provide a qualitative comparison between TD and scMoMaT. However, because UMAP visualization alone is insufficient for benchmarking, the comparison was primarily evaluated using the quantitative metrics described above.

**Fig. 11.**
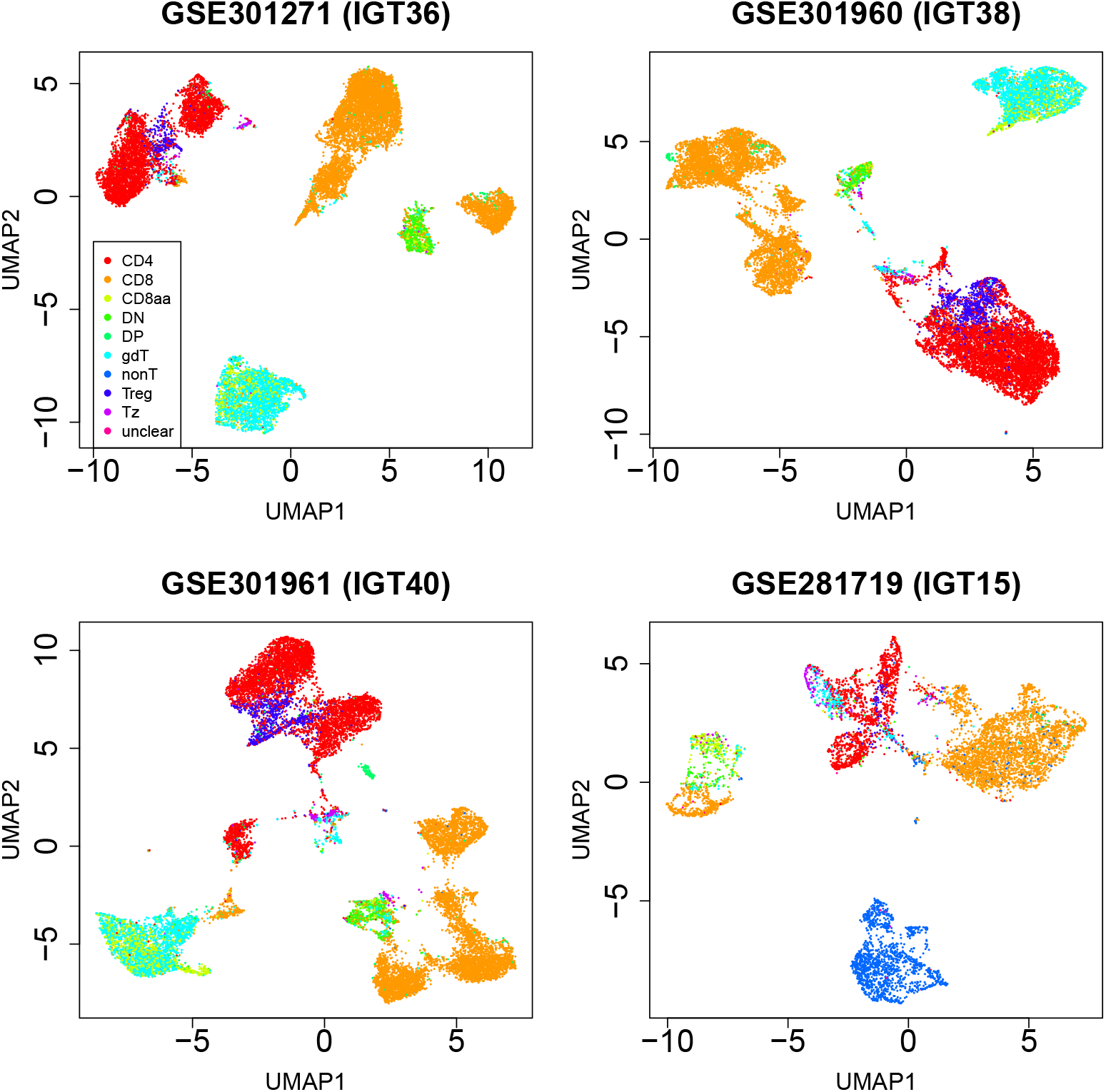
UMAP embedding by scMoMaT. The coloring indicates level 1 classification of metadata (cell types)

Compared with scMoMaT, TD-based UFE required less computational time and memory under the tested settings (Table 11). For the three largest datasets, TD was approximately 3.8- to 7.0-fold faster and used about one-sixth of the memory required by scMoMaT. Together with the quantitative cell-type consistency metrics, these results suggest that TD-based UFE can serve as a computationally lightweight factorization-based reference method for paired RNA/ADT CITE-seq data. However, this comparison should be interpreted cautiously because scMoMaT was originally developed for mosaic multi-omics integration, whereas the present study analyzed fully paired RNA/ADT datasets. Thus, the comparison supports the practical utility of TD-based UFE under the tested setting, but does not establish general superiority over all multimodal integration methods.

**Table 11.** Comparison of required CPU time and memory size between TD and scMoMaT. The SPEC of computer:Intel(R) Xeon(R) CPU E5-2643 v4 3.40GHz, 252 GB, 12 CPU. CPU time is counted as single CPU performance base.

| Data set | CPU time |  | Memory |  |
| --- | --- | --- | --- | --- |
|  | TD | scMoMaT | TD | scMoMaT |
| GSE301271 | 4 hours | 15 hours | 1GB | 6.3 GB |
| GSE301960 | 2 hours | 14 hours | 1GB | 6.2 GB |
| GSE301961 | 2 hours | 14 hours | 1GB | 6.1 GB |

The quantitative benchmarking clarifies the appropriate scope of TD-based UFE. TD-based UFE is not a replacement for WNN or ADT-only cell-type analysis.

Its value is that it provides a lightweight tensor-decomposition-based framework for paired RNA/ADT CITE-seq data, linking cell-side representation and post hoc unsupervised gene selection without explicit modality-weight tuning or prior HVG filtering. ADT-only embeddings and Seurat WNN graphs are highly effective for cell-type neighborhood construction, whereas TD-based UFE provides an interpretable factorization-based mechanism for connecting RNA/ADT-integrated cell representations with associated genes.

totalVI was not retrained from raw counts for all datasets because no GPU resources were available and CPU training was prohibitively slow. Instead, we used publicly available totalVI embeddings when available, and clearly treated them as reference embeddings rather than fully controlled reimplementations.

One possible concern is that the selected set of 6,707 genes is relatively large, and that broad T-cell- or lymph-node-related enrichment may be expected simply because the dataset is T-cell-focused. To address this concern, we compared the TD-selected genes with the same number of highly variable genes (HVGs). The HVG-based gene set was also subjected to Enrichr analysis, and the results are summarized in Tables S8–S11. Compared with the HVG reference, TD-selected genes showed stronger or more specific enrichment in several biologically relevant categories. In particular, TD-selected genes showed smaller adjusted P-values and a larger number of CD4/CD8- and lymph-node-related enriched terms in the tested Enrichr libraries. For the Tabula Sapiens category, the HVG reference produced only a small number of significant enriched terms, whereas TD-selected genes showed clearer lymph-node-related enrichment. These results suggest that the enrichment observed for TD-selected genes was not simply a consequence of selecting a large number of genes.

We also compared enrichment results between TD-selected genes and RNA-only-selected genes. Cell-type-related enrichment categories are summarized in Tables 4, 5, 6, and 7, KEGG pathway enrichment is summarized in Table 8, and GO BP, GO CC, and GO MF enrichment results are summarized in Tables S12–S14, respectively. In the cell-type-related enrichment categories, TD-selected and RNA-only-selected genes showed broadly comparable enrichment significance, although several terms differed between the two gene sets. In contrast, clearer differences were observed in KEGG and GO enrichment analyses: TD-selected genes showed stronger or more specific enrichment than the RNA-only reference gene set in several KEGG pathways and GO BP/CC/MF categories. These comparisons support the interpretation that TD-selected genes capture biological structure associated with the RNA/ADT-integrated representation, while the enrichment results should still be regarded as exploratory biological plausibility checks rather than definitive validation.

The comparison with scMoMaT should also be interpreted cautiously. scMoMaT was originally developed for mosaic multi-omics integration, whereas the present study analyzed fully paired RNA/ADT datasets. MOFA+ would be another relevant factorization-based reference; however, in large single-cell applications it is commonly used after feature filtering for computational and statistical reasons. Because one aim of TD-based UFE is to avoid prior HVG-based gene filtering and instead perform post hoc unsupervised gene selection, we did not treat MOFA+ as the primary comparator in this study. Future work should evaluate TD-based UFE against MOFA+ under matched feature-selection settings.

The product-form tensor construction is not an ad hoc choice introduced only for the present CITE-seq analysis. It follows previous TD-based UFE applications in which two omics profiles measured on the same samples were combined by an outer-product representation [13]: gene expression and methylation [20], mRNA and miRNA expression [21], gene expression and transcriptome [22], and protein-protein interaction and gene expression [23]. These previous applications motivated us to extend the same construction to paired RNA/ADT CITE-seq data. At the same time, this tensor should be interpreted as a representation of cell-wise co-occurrence between RNA and ADT measurements, not as a direct mechanistic model of gene–protein regulation. Because the numerical values of the tensor depend on the input RNA and ADT matrices, pre-processing choices such as RNA normalization and ADT transformation may affect the decomposition. Thus, the present study evaluates this established TD-based construction in the CITE-seq setting, while robustness under alternative preprocessing schemes remains to be examined.

Several limitations of the present study should be emphasized. First, although the proposed workflow was evaluated across multiple datasets, we did not perform a systematic sensitivity analysis covering the number of cell-mode singular vectors, gene-selection thresholds, UMAP parameters and random seeds, and alternative RNA and ADT normalization schemes. The robustness of the results to these analytical choices should therefore be evaluated more comprehensively in future studies. Second, the present interpretation is group-oriented: TD-based unsupervised FE identifies genes associated with the RNA/ADT-integrated representation, but we did not establish one-to-one relationships between individual tensor factors, specific ADT markers, and RNA programs. Factor-specific analyses linking gene and protein loadings to cell types and biological programs constitute an important direction for future work. Third, the enrichment analyses were performed using the default Enrichr background and should be regarded as exploratory biological plausibility checks rather than definitive functional validation. These limitations do not affect the quantitative comparisons presented here, but they define the appropriate scope of the present study and motivate further investigation.

## 5 Conclusion

In conclusion, TD-based UFE provides a tensor-decomposition-based framework for paired RNA/ADT CITE-seq analysis without explicit RNA/ADT modality-weight tuning or prior HVG-based gene filtering. The method yields cell-mode singular vectors and enables post hoc unsupervised gene selection from the same integrated tensor structure. Quantitative comparisons showed that TD-derived representations were competitive with scMoMaT, a related factorization-based reference, but did not outperform ADT-only embeddings or Seurat WNN graphs in cell-type neighborhood consistency. Thus, TD-based UFE should not be regarded as a replacement for WNN, ADT-only, or marker-based cell-type analysis. Rather, its main contribution is to provide an interpretable tensor-based feature-extraction framework that links RNA/ADT-integrated cell representations with post hoc gene selection. Future work should evaluate its robustness under alternative normalization schemes, matched MOFA+ settings, and incomplete or mosaic multi-omics designs.

## Supplementary information

Supplementary file 1 [xlsx] : List of 6707 genes selected by TD and associated enrichment analysis

Supplementary file 2 [xlsx] : List of 6707 genes selected by HVG and associated enrichment analysis

Supplementary file 3 [xlsx] : List of 6707 genes selected by RNA only and associated enrichment analysis

Supplementary file 4 [xlsx] : Supplementary Tables

## Declarations

- Funding: The project was funded by KAU Endowment (WAQF) at King Abdulaziz University, Jeddah, Saudi Arabia. The authors, therefore, acknowledge and thank WAQF and the Deanship of Scientific Research (DSR) for technical and financial support.
- Conflict of interest/Competing interests : The authors declare no conflict of interest.
- Ethics approval and consent to participate : NA
- Consent for publication : NA
- Data availability : All data can be accessed through GSE301271, GSE301960, GSE301961, GSE281719, and GSE283941.
- Materials availability : NA
- Code availability : Sample code is in https://github.com/tagtag/TDbasedUFECITE-seq
- Author contributions : Y. H. T. planned the study and performed the analyses. Y.-H.T. and T.T. evaluated the results and wrote and reviewed the manuscript. Conceptualization, data curation, and analysis were performed by Y. H. T. All the authors have read and agreed to the published version of the manuscript.

## Appendix A TD using columnar-blocked matrices

### A.1 Implicit tensor model

Given sparse matrices

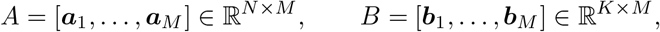

defines the third-order tensor as

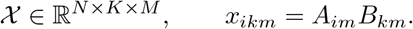

Similarly, the *m*th frontal slice is:

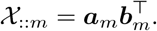

Seeking truncated HOSVD

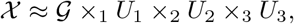

where

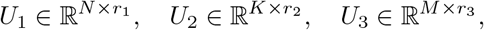

have orthonormal columns and

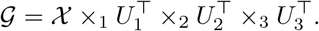

### A.2 Mode-wise Gram matrices

The HOSVD factor matrices are obtained from the leading eigenvectors of the

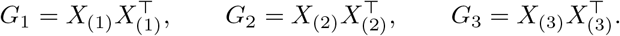

For the implicit tensor above, these matrices can be written as

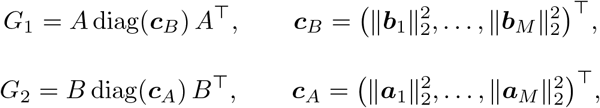

and

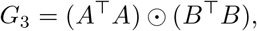

where ⊙ denotes the Hadamard product,

#### Algorithm 1

Truncated HOSVD for the implicit tensor *x*_*ikm*_ = *A*_*im*_*B*_*km*_

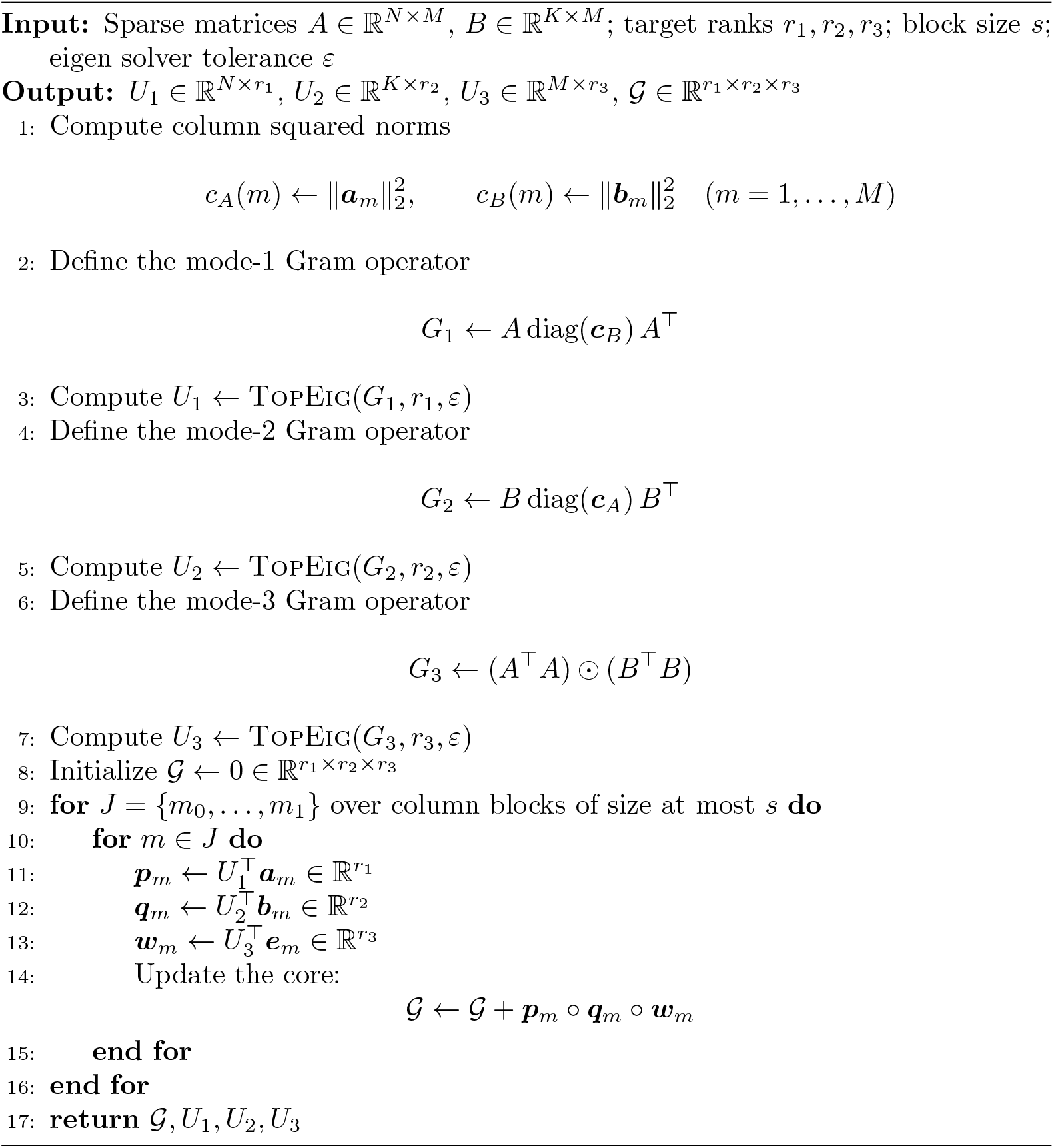

### A.3 Equivalent expression for the core

The above blockwise update is equivalent to

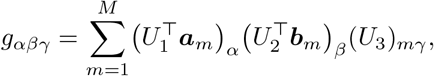

or, equivalently,

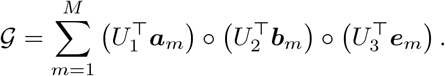

### A.4 Remark

During the implementation of large sparse matrices, operators

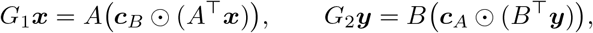

and

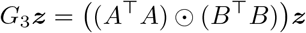

are typically applied implicitly inside an iterative eigensolver, rather than densely forming all Gram matrices.

## References

[1] Hao, Y., et al.: Integrated analysis of multimodal single-cell data. Cell 184(13), 3573–358729 (2021)

[2] Kim, H.J., et al.: Citefuse enables multi-modal analysis of CITE-seq data. Bioinformatics 36(14), 4137–4143 (2020) 10.1093/bioinformatics/btaa282

[3] Quinn, T.P., Richardson, M.F., Lovell, D., Crowley, T.M.: propr: An r-package for identifying proportionally abundant features using compositional data analysis. Scientific Reports 7(1), 16252 (2017) 10.1038/s41598-017-16520-0

[4] Gayoso, A., et al.: Joint probabilistic modeling of single-cell multi-omic data with totalVI. Nature Methods 18(3), 272–282 (2021) 10.1038/s41592-020-01050-x

[5] Lakkis, J., Schroeder, A., Su, K., Lee, M.Y.Y., Bashore, A.C., Reilly, M.P., Li, M.: A multi-use deep learning method for CITE-seq and single-cell rna-seq data integration with cell surface protein prediction and imputation. Nature Machine Intelligence 4(11), 940–952 (2022) 10.1038/s42256-022-00545-w

[6] Yuan, M., Chen, L., Deng, M.: Clustering cite-seq data with a canonical correlation-based deep learning method. Frontiers in Genetics Volume 13 - 2022 (2022) 10.3389/fgene.2022.977968

[7] Lin, X., Tian, T., Wei, Z., Hakonarson, H.: Clustering of single-cell multi-omics data with a multimodal deep learning method. Nature Communications 13(1), 7705 (2022) 10.1038/s41467-022-35031-9

[8] Argelaguet, R., Arnol, D., Bredikhin, D., Deloro, Y., Velten, B., Marioni, J.C., Stegle, O.: MOFA+: a statistical framework for comprehensive integration of multi-modal single-cell data. Genome Biology 21(1), 111 (2020)

[9] Zhang, Z., Sun, H., Mariappan, R., Chen, X., Chen, X., Jain, M.S., Efremova, M., Teichmann, S.A., Rajan, V., Zhang, X.: scMoMaT jointly performs single cell mosaic integration and multi-modal bio-marker detection. Nature Communications 14(1), 384 (2023) 10.1038/s41467-023-36066-2

[10] He, P., et al.: A deep probabilistic framework for mosaic integration and knowledge transfer of single-cell multimodal data. Nature Biotechnology (2024) 10.1038/s41587-023-02040-y

[11] Ghazanfar, S., Guibentif, C., Marioni, J.C.: Stabilized mosaic single-cell data integration using unshared features. Nature Biotechnology 42(2), 284–292 (2024) 10.1038/s41587-023-01766-z

[12] Hao, Y., Stuart, T., Kowalski, M., Choudhary, S., Hoffman, P., Hartman, A., Srivastava, A., Molla, G., Madad, S., Fernandez-Granda, C., Satija, R.: Dictionary learning for integrative, multimodal and scalable single-cell analysis. Nature Biotechnology 42(2), 293–304 (2024) 10.1038/s41587-023-01767-y

[13] Taguchi, Y.-h.: Unsupervised Feature Extraction Applied to Bioinformatics: A PCA Based and TD Based Approach, 2nd edn. Unsupervised and Semi-Supervised Learning. Springer, Switzland (2024). 10.1007/978-3-031-60982-4. 10.1007/978-3-031-60982-4

[14] Zemmour, D., Goldrath, A., Kronenberg, M., Kang, J., Benoist, C.: The immgen consortium opensource t cell project. Nature Immunology 23(5), 643–644 (2022) 10.1038/s41590-022-01197-z

[15] Bates, D., Maechler, M., Jagan, M.: Matrix: Sparse and Dense Matrix Classes and Methods. (2025). 10.32614/CRAN.package.Matrix . R package version 1. 7–3. https://CRAN.R-project.org/package=Matrix

[16] Konopka, T.: Umap: Uniform Manifold Approximation and Projection. (2023). 10.32614/CRAN.package.umap . R package version 0.2.10.0. https://CRAN.R-project.org/package=umap

[17] Xie, Z., Bailey, A., Kuleshov, M.V., Clarke, D.J.B., Evangelista, J.E., Jenkins, S.L., Lachmann, A., Wojciechowicz, M.L., Kropiwnicki, E., Jagodnik, K.M., Jeon, M., Ma’ayan, A.: Gene set knowledge discovery with enrichr. Current Protocols 1(3), 90 (2021) 10.1002/cpz1.90 https://currentprotocols.onlinelibrary.wiley.com/doi/pdf/10.1002/cpz1.90

[18] Kanehisa, M., Furumichi, M., Sato, Y., Matsuura, Y., Ishiguro-Watanabe, M.: KEGG: biological systems database as a model of the real world. Nucleic Acids Research 53(D1), 672–677 (2025) 10.1093/nar/gkae909

[19] Taguchi, Y.-h., Turki, T.: Tensor-decomposition-based unsupervised feature extraction in single-cell multiomics data analysis. Genes 12(9) (2021) 10.3390/genes12091442

[20] Y, q.H.T.: Tensor decomposition-based and principal-component-analysis-based unsupervised feature extraction applied to the gene expression and methylation profiles in the brains of social insects with multiple castes. BMC Bioinformatics 19(Suppl 4), 99 (2018) 10.1186/s12859-018-2068-7

[21] Ng, K.-L., Y, q.H.T.: Identification of miRNA signatures for kidney renal clear cell carcinoma using the tensor-decomposition method. Scientific Reports 10(1), 15149 (2020) 10.1038/s41598-020-71997-6

[22] Roy, S.S., Y, q.H.T.: Identification of genes associated with altered gene expression and m6a profiles during hypoxia using tensor decomposition based unsupervised feature extraction. Scientific Reports 11(1), 8909 (2021) 10.1038/s41598-021-87779-7

[23] Taguchi, Y.-H., Turki, T.: Integrated analysis of gene expression and protein–protein interaction with tensor decomposition. Mathematics 11(17) (2023) 10.3390/math11173655

