## Supplementary material for "Interpretable Integration of CITE-seq RNA and ADT Profiles Without Explicit Modality-Weight Tuning via Tensor Decomposition-Based Unsupervised Feature Extraction": Supplementary_file_4.pdf

Table S1: Cell type specific accuracy

| cell type | TD accuracy | scMoMaT accuracy | difference |
|---|---|---|---|
| | | GSE301271 | |
| CD4 | 0.989 | 0.927 | +0.061 |
| CD8 | 0.968 | 0.959 | +0.009 |
| CD8aa | 0.091 | 0.127 | -0.036 |
| DN | 0.841 | 0.772 | +0.069 |
| DP | 0.196 | 0.056 | +0.140 |
| gdT | 0.897 | 0.856 | +0.041 |
| nonT | 0.000 | 0.000 | +0.000 |
| Treg | 0.325 | 0.264 | +0.061 |
| Tz | 0.472 | 0.304 | +0.168 |
| unclear | 0.000 | 0.000 | +0.000 |
| | | GSE301960 | |
| CD4 | 0.966 | 0.956 | +0.009 |
| CD8 | 0.964 | 0.948 | +0.017 |
| CD8aa | 0.209 | 0.231 | -0.023 |
| DN | 0.789 | 0.026 | +0.764 |
| DP | 0.277 | 0.012 | +0.265 |
| gdT | 0.883 | 0.813 | +0.070 |
| nonT | 0.095 | 0.190 | -0.095 |
| Treg | 0.541 | 0.582 | -0.041 |
| Tz | 0.142 | 0.027 | +0.115 |
| unclear | 0.000 | 0.000 | +0.000 |
| | | GSE301961 | |
| CD4 | 0.965 | 0.958 | +0.007 |
| CD8 | 0.963 | 0.955 | +0.008 |
| CD8aa | 0.371 | 0.367 | +0.005 |
| DN | 0.778 | 0.628 | +0.150 |
| DP | 0.284 | 0.092 | +0.191 |
| gdT | 0.676 | 0.724 | -0.048 |
| nonT | 0.000 | 0.107 | -0.107 |
| Treg | 0.439 | 0.378 | +0.061 |
| Tz | 0.558 | 0.550 | +0.008 |
| unclear | 0.000 | 0.000 | +0.000 |
| | | GSE281719 | |
| CD4 | 0.950 | 0.798 | +0.152 |
| CD8 | 0.977 | 0.970 | +0.007 |
| CD8aa | 0.604 | 0.532 | +0.072 |
| DN | 0.076 | 0.102 | -0.025 |
| DP | 0.000 | 0.000 | 0.000 |
| gdT | 0.291 | 0.384 | -0.094 |
| nonT | 0.928 | 0.911 | +0.016 |
| Treg | 0.000 | 0.000 | 0.000 |
| Tz | 0.021 | 0.264 | -0.244 |
| unclear | 0.000 | 0.000 | 0.000 |

Table S2: Various measures about the consistency between TD embedding / Seurat WNN graph and cell types

| Dataset | Index | TD | WNN | Interpretation |
|---|---|---|---|---|
| GSE301271 | kNN purity / graph kNN purity | 0.8396 | 0.8902 | TD < WNN |
| GSE301271 | macro kNN purity / macro graph kNN purity | 0.4462 | 0.5675 | TD < WNN |
| GSE301271 | kNN classification accuracy / graph kNN classification accuracy | 0.8876 | 0.9214 | TD < WNN |
| GSE301271 | macro kNN accuracy / macro graph kNN accuracy | 0.4778 | 0.5922 | TD < WNN |
| GSE301960 | kNN purity / graph kNN purity | 0.8230 | 0.9020 | TD < WNN |
| GSE301960 | macro kNN purity / macro graph kNN purity | 0.4493 | 0.6244 | TD < WNN |
| GSE301960 | kNN classification accuracy / graph kNN classification accuracy | 0.8743 | 0.9330 | TD < WNN |
| GSE301960 | macro kNN accuracy / macro graph kNN accuracy | 0.4866 | 0.6857 | TD < WNN |
| GSE301961 | kNN purity / graph kNN purity | 0.7900 | 0.8750 | TD < WNN |
| GSE301961 | macro kNN purity / macro graph kNN purity | 0.4500 | 0.6394 | TD < WNN |
| GSE301961 | kNN classification accuracy / graph kNN classification accuracy | 0.8350 | 0.9209 | TD < WNN |
| GSE301961 | macro kNN accuracy / macro graph kNN accuracy | 0.5034 | 0.6935 | TD < WNN |
| GSE281719 | kNN purity / graph kNN purity | 0.8040 | 0.8984 | TD < WNN |
| GSE281719 | macro kNN purity / macro graph kNN purity | 0.3771 | 0.6159 | TD < WNN |
| GSE281719 | kNN classification accuracy / graph kNN classification accuracy | 0.8592 | 0.9294 | TD < WNN |
| GSE281719 | macro kNN accuracy / macro graph kNN accuracy | 0.3846 | 0.6312 | TD < WNN |

Table S3: Cell type-specific kNN classification accuracy of TD and graph kNN classification accuracy of Seurat WNN

| Dataset | cell type | TD accuracy | WNN accuracy | Difference |
|---|---|---|---|---|
| GSE301271 | CD4 | 0.989 | 0.977 | +0.012 |
| GSE301271 | CD8 | 0.968 | 0.971 | -0.003 |
| GSE301271 | CD8aa | 0.091 | 0.364 | -0.273 |
| GSE301271 | DN | 0.841 | 0.895 | -0.054 |
| GSE301271 | DP | 0.196 | 0.189 | +0.007 |
| GSE301271 | gdT | 0.897 | 0.937 | -0.040 |
| GSE301271 | nonT | 0.000 | 0.000 | +0.000 |
| GSE301271 | Treg | 0.325 | 0.838 | -0.513 |
| GSE301271 | Tz | 0.472 | 0.752 | -0.280 |
| GSE301271 | unclear | 0.000 | 0.000 | +0.000 |
| GSE301960 | CD4 | 0.966 | 0.982 | -0.017 |
| GSE301960 | CD8 | 0.964 | 0.965 | -0.000 |
| GSE301960 | CD8aa | 0.209 | 0.543 | -0.334 |
| GSE301960 | DN | 0.789 | 0.789 | +0.000 |
| GSE301960 | DP | 0.277 | 0.283 | -0.006 |
| GSE301960 | gdT | 0.883 | 0.948 | -0.065 |
| GSE301960 | nonT | 0.095 | 0.857 | -0.762 |
| GSE301960 | Treg | 0.541 | 0.935 | -0.394 |
| GSE301960 | Tz | 0.142 | 0.554 | -0.412 |
| GSE301960 | unclear | 0.000 | 0.000 | +0.000 |
| GSE301961 | CD4 | 0.965 | 0.975 | -0.010 |
| GSE301961 | CD8 | 0.963 | 0.954 | +0.009 |
| GSE301961 | CD8aa | 0.371 | 0.803 | -0.431 |
| GSE301961 | DN | 0.778 | 0.739 | +0.039 |
| GSE301961 | DP | 0.284 | 0.257 | +0.026 |
| GSE301961 | gdT | 0.676 | 0.911 | -0.236 |
| GSE301961 | nonT | 0.000 | 0.643 | -0.643 |
| GSE301961 | Treg | 0.439 | 0.898 | -0.460 |
| GSE301961 | Tz | 0.558 | 0.754 | -0.196 |
| GSE301961 | unclear | 0.000 | 0.000 | +0.000 |
| GSE281719 | CD4 | 0.950 | 0.941 | +0.009 |
| GSE281719 | CD8 | 0.977 | 0.982 | -0.004 |
| GSE281719 | CD8aa | 0.604 | 0.838 | -0.234 |
| GSE281719 | DN | 0.076 | 0.513 | -0.436 |
| GSE281719 | DP | 0.000 | 0.000 | +0.000 |
| GSE281719 | gdT | 0.291 | 0.681 | -0.391 |
| GSE281719 | nonT | 0.928 | 0.987 | -0.059 |
| GSE281719 | Treg | 0.000 | 0.660 | -0.660 |
| GSE281719 | Tz | 0.021 | 0.710 | -0.689 |
| GSE281719 | unclear | 0.000 | 0.000 | +0.000 |

Table S4: Various measures about the consistency between TD and RNA-only embedding and cell types

| Dataset | Index | TD | RNA | Interpretation |
|---|---|---|---|---|
| GSE301271 | kNN purity | 0.8396 | 0.8187 | TD > RNA |
| GSE301271 | macro kNN purity | 0.4462 | 0.4366 | TD > RNA |
| GSE301271 | kNN classification accuracy | 0.8876 | 0.8775 | TD > RNA |
| GSE301271 | macro kNN accuracy | 0.4778 | 0.4778 | TD $\simeq$ RNA |
| GSE301271 | silhouette | -0.1825 | 0.0088 | TD < RNA |
| GSE301960 | kNN purity | 0.8230 | 0.8064 | TD > RNA |
| GSE301960 | macro kNN purity | 0.4493 | 0.4582 | TD < RNA |
| GSE301960 | kNN classification accuracy | 0.8743 | 0.8699 | TD > RNA |
| GSE301960 | macro kNN accuracy | 0.4866 | 0.4966 | TD < RNA |
| GSE301960 | silhouette | -0.1764 | 0.0445 | TD < RNA |
| GSE301961 | kNN purity | 0.7900 | 0.7218 | TD > RNA |
| GSE301961 | macro kNN purity | 0.4500 | 0.4141 | TD > RNA |
| GSE301961 | kNN classification accuracy | 0.8350 | 0.8006 | TD > RNA |
| GSE301961 | macro kNN accuracy | 0.5034 | 0.4626 | TD > RNA |
| GSE301961 | silhouette | -0.1545 | -0.0622 | TD < RNA |
| GSE281719 | kNN purity | 0.8040 | 0.8487 | TD < RNA |
| GSE281719 | macro kNN purity | 0.3771 | 0.4967 | TD < RNA |
| GSE281719 | kNN classification accuracy | 0.8592 | 0.8915 | TD < RNA |
| GSE281719 | macro kNN accuracy | 0.3846 | 0.5105 | TD < RNA |
| GSE281719 | silhouette | -0.0154 | 0.0390 | TD < RNA |

Table S5: Cell type-specific kNN classification accuracy of TD and RNA-only embedding

| Dataset | cell type | TD accuracy | RNA accuracy | Difference |
|---|---|---|---|---|
| GSE301271 | CD4 | 0.989 | 0.955 | +0.034 |
| GSE301271 | CD8 | 0.968 | 0.957 | +0.011 |
| GSE301271 | CD8aa | 0.091 | 0.173 | -0.082 |
| GSE301271 | DN | 0.841 | 0.869 | -0.028 |
| GSE301271 | DP | 0.196 | 0.000 | +0.196 |
| GSE301271 | gdT | 0.897 | 0.896 | +0.001 |
| GSE301271 | nonT | 0.000 | 0.000 | +0.000 |
| GSE301271 | Treg | 0.325 | 0.305 | +0.020 |
| GSE301271 | Tz | 0.472 | 0.624 | -0.152 |
| GSE301271 | unclear | 0.000 | 0.000 | +0.000 |
| GSE301960 | CD4 | 0.966 | 0.947 | +0.019 |
| GSE301960 | CD8 | 0.964 | 0.931 | +0.034 |
| GSE301960 | CD8aa | 0.209 | 0.254 | -0.046 |
| GSE301960 | DN | 0.789 | 0.718 | +0.071 |
| GSE301960 | DP | 0.277 | 0.024 | +0.253 |
| GSE301960 | gdT | 0.883 | 0.879 | +0.004 |
| GSE301960 | nonT | 0.095 | 0.048 | +0.048 |
| GSE301960 | Treg | 0.541 | 0.794 | -0.253 |
| GSE301960 | Tz | 0.142 | 0.372 | -0.230 |
| GSE301960 | unclear | 0.000 | 0.000 | +0.000 |
| GSE301961 | CD4 | 0.965 | 0.916 | +0.050 |
| GSE301961 | CD8 | 0.963 | 0.900 | +0.064 |
| GSE301961 | CD8aa | 0.371 | 0.444 | -0.073 |
| GSE301961 | DN | 0.778 | 0.476 | +0.302 |
| GSE301961 | DP | 0.284 | 0.026 | +0.257 |
| GSE301961 | gdT | 0.676 | 0.692 | -0.016 |
| GSE301961 | nonT | 0.000 | 0.000 | +0.000 |
| GSE301961 | Treg | 0.439 | 0.657 | -0.218 |
| GSE301961 | Tz | 0.558 | 0.515 | +0.042 |
| GSE301961 | unclear | 0.000 | 0.000 | +0.000 |
| GSE281719 | CD4 | 0.950 | 0.904 | +0.046 |
| GSE281719 | CD8 | 0.977 | 0.970 | +0.008 |
| GSE281719 | CD8aa | 0.604 | 0.833 | -0.230 |
| GSE281719 | DN | 0.076 | 0.363 | -0.287 |
| GSE281719 | DP | 0.000 | 0.000 | +0.000 |
| GSE281719 | gdT | 0.291 | 0.603 | -0.312 |
| GSE281719 | nonT | 0.928 | 0.965 | -0.038 |
| GSE281719 | Treg | 0.000 | 0.000 | +0.000 |
| GSE281719 | Tz | 0.021 | 0.466 | -0.446 |
| GSE281719 | unclear | 0.000 | 0.000 | +0.000 |

Table S6: Various measures about the consistency between TD and ADT-only embedding and cell types

| Dataset | Index | TD | ADT | Interpretation |
|---|---|---|---|---|
| GSE301271 | kNN purity | 0.8396 | 0.8381 | TD $\simeq$ ADT |
| GSE301271 | macro kNN purity | 0.4462 | 0.4690 | TD $<$ ADT |
| GSE301271 | kNN classification accuracy | 0.8876 | 0.9130 | TD $<$ ADT |
| GSE301271 | macro kNN accuracy | 0.4778 | 0.5365 | TD $<$ ADT |
| GSE301271 | silhouette | -0.1825 | -0.0168 | TD $<$ ADT |
| GSE301960 | kNN purity | 0.8230 | 0.8410 | TD $<$ ADT |
| GSE301960 | macro kNN purity | 0.4493 | 0.5044 | TD $<$ ADT |
| GSE301960 | kNN classification accuracy | 0.8743 | 0.9147 | TD $<$ ADT |
| GSE301960 | macro kNN accuracy | 0.4866 | 0.5734 | TD $<$ ADT |
| GSE301960 | silhouette | -0.1764 | 0.0166 | TD $<$ ADT |
| GSE301961 | kNN purity | 0.7900 | 0.8300 | TD $<$ ADT |
| GSE301961 | macro kNN purity | 0.4500 | 0.5038 | TD $<$ ADT |
| GSE301961 | kNN classification accuracy | 0.8350 | 0.9030 | TD $<$ ADT |
| GSE301961 | macro kNN accuracy | 0.5034 | 0.5524 | TD $<$ ADT |
| GSE301961 | silhouette | -0.1545 | 0.0195 | TD $<$ ADT |
| GSE281719 | kNN purity | 0.8040 | 0.8459 | TD $<$ ADT |
| GSE281719 | macro kNN purity | 0.3771 | 0.5107 | TD $<$ ADT |
| GSE281719 | kNN classification accuracy | 0.8592 | 0.9064 | TD $<$ ADT |
| GSE281719 | macro kNN accuracy | 0.3846 | 0.5608 | TD $<$ ADT |
| GSE281719 | silhouette | -0.0154 | 0.0212 | TD $<$ ADT |

Table S7: Cell type-specific kNN classification accuracy of TD and ADT-only embedding

| Dataset | cell type | TD accuracy | ADT accuracy | Difference |
|---|---|---|---|---|
| GSE301271 | CD4 | 0.989 | 0.964 | +0.025 |
| GSE301271 | CD8 | 0.968 | 0.979 | -0.011 |
| GSE301271 | CD8aa | 0.091 | 0.527 | -0.436 |
| GSE301271 | DN | 0.841 | 0.792 | +0.049 |
| GSE301271 | DP | 0.196 | 0.007 | +0.189 |
| GSE301271 | gdT | 0.897 | 0.888 | +0.009 |
| GSE301271 | nonT | 0.000 | 0.000 | +0.000 |
| GSE301271 | Treg | 0.325 | 0.728 | -0.404 |
| GSE301271 | Tz | 0.472 | 0.480 | -0.008 |
| GSE301271 | unclear | 0.000 | 0.000 | +0.000 |
| GSE301960 | CD4 | 0.966 | 0.980 | -0.014 |
| GSE301960 | CD8 | 0.964 | 0.975 | -0.010 |
| GSE301960 | CD8aa | 0.209 | 0.584 | -0.375 |
| GSE301960 | DN | 0.789 | 0.650 | +0.140 |
| GSE301960 | DP | 0.277 | 0.018 | +0.259 |
| GSE301960 | gdT | 0.883 | 0.910 | -0.027 |
| GSE301960 | nonT | 0.095 | 0.714 | -0.619 |
| GSE301960 | Treg | 0.541 | 0.788 | -0.247 |
| GSE301960 | Tz | 0.142 | 0.115 | +0.027 |
| GSE301960 | unclear | 0.000 | 0.000 | +0.000 |
| GSE301961 | CD4 | 0.965 | 0.976 | -0.010 |
| GSE301961 | CD8 | 0.963 | 0.974 | -0.011 |
| GSE301961 | CD8aa | 0.371 | 0.804 | -0.433 |
| GSE301961 | DN | 0.778 | 0.607 | +0.172 |
| GSE301961 | DP | 0.284 | 0.026 | +0.257 |
| GSE301961 | gdT | 0.676 | 0.858 | -0.182 |
| GSE301961 | nonT | 0.000 | 0.036 | -0.036 |
| GSE301961 | Treg | 0.439 | 0.767 | -0.328 |
| GSE301961 | Tz | 0.558 | 0.477 | +0.081 |
| GSE301961 | unclear | 0.000 | 0.000 | +0.000 |
| GSE281719 | CD4 | 0.950 | 0.904 | +0.046 |
| GSE281719 | CD8 | 0.977 | 0.994 | -0.017 |
| GSE281719 | CD8aa | 0.604 | 0.739 | -0.135 |
| GSE281719 | DN | 0.076 | 0.338 | -0.261 |
| GSE281719 | DP | 0.000 | 0.000 | +0.000 |
| GSE281719 | gdT | 0.291 | 0.753 | -0.462 |
| GSE281719 | nonT | 0.928 | 0.939 | -0.011 |
| GSE281719 | Treg | 0.000 | 0.476 | -0.476 |
| GSE281719 | Tz | 0.021 | 0.466 | -0.446 |
| GSE281719 | unclear | 0.000 | 0.000 | +0.000 |

Table S8: Top 10 enriched terms for "Azimuth Cell Types 2021" category in Enrichr for HVG. Full list is available in the Supplementary Materials.

| Term | Overlap | $P$-value | Adjusted $P$-value |
|---|---|---|---|
| Innate Lymphoid Cell CL0001065 | 13/14 | $6.49 \times 10^{-6}$ | $2.16 \times 10^{-3}$ |
| Cycling CL0008024 | 10/10 | $1.79 \times 10^{-5}$ | $2.98 \times 10^{-3}$ |
| Proliferating Natural Killer CL0000623 | 12/14 | $8.74 \times 10^{-5}$ | $9.70 \times 10^{-3}$ |
| Natural Killer T CL0000814 | 10/11 | $1.37 \times 10^{-4}$ | $1.03 \times 10^{-2}$ |
| Plasmablast CL0000980 | 14/18 | $1.55 \times 10^{-4}$ | $1.03 \times 10^{-2}$ |
| CD8+ Naive T CL0000900 | 17/24 | $2.08 \times 10^{-4}$ | $1.04 \times 10^{-2}$ |
| CD4+ Proliferating T CL0000896 | 11/13 | $2.25 \times 10^{-4}$ | $1.04 \times 10^{-2}$ |
| CD4+ Central Memory T 3 CL0000897 | 9/10 | $3.73 \times 10^{-4}$ | $1.04 \times 10^{-2}$ |
| Intermediate B Cell CL0000785 | 9/10 | $3.73 \times 10^{-4}$ | $1.04 \times 10^{-2}$ |
| Other T Cell CL0002419 | 9/10 | $3.73 \times 10^{-4}$ | $1.04 \times 10^{-2}$ |

Table S9: Top 10 enriched terms for "PanglaoDB Augmented 2021" category in Enrichr for HVG. Full list is available in Supplementary material.

| Term | Overlap | $P$-value | Adjusted $P$-value |
|---|---|---|---|
| T Helper Cells | 100/148 | $2.35 \times 10^{-17}$ | $4.19 \times 10^{-15}$ |
| T Cells | 102/165 | $8.51 \times 10^{-14}$ | $7.57 \times 10^{-12}$ |
| NK Cells | 96/157 | $1.16 \times 10^{-12}$ | $6.89 \times 10^{-11}$ |
| Macrophages | 115/204 | $1.53 \times 10^{-11}$ | $6.80 \times 10^{-10}$ |
| Natural Killer T Cells | 75/118 | $2.30 \times 10^{-11}$ | $8.18 \times 10^{-10}$ |
| T Regulatory Cells | 74/117 | $4.40 \times 10^{-11}$ | $1.31 \times 10^{-9}$ |
| Monocytes | 96/176 | $7.28 \times 10^{-9}$ | $1.85 \times 10^{-7}$ |
| Pluripotent Stem Cells | 65/112 | $8.66 \times 10^{-8}$ | $1.93 \times 10^{-6}$ |
| T Cytotoxic Cells | 60/102 | $1.39 \times 10^{-7}$ | $2.74 \times 10^{-6}$ |
| Microglia | 84/158 | $2.75 \times 10^{-7}$ | $4.90 \times 10^{-6}$ |

Table S10: Top 10 enriched terms for the "CellMarker Augmented 2021" category in Enrichr for HVG. Full list is available in the Supplementary Materials.

| Term | Overlap | $P$-value | Adjusted $P$-value |
|---|---|---|---|
| Oxyntic Stem cell:Stomach | 94/99 | $1.57 \times 10^{-38}$ | $5.70 \times 10^{-36}$ |
| Oxyntic Progenitor cell:Stomach | 94/99 | $1.57 \times 10^{-38}$ | $5.70 \times 10^{-36}$ |
| Progenitor cell:Stomach | 94/99 | $1.57 \times 10^{-38}$ | $5.70 \times 10^{-36}$ |
| Vascular Stem cell:Blood | 94/100 | $1.75 \times 10^{-37}$ | $3.83 \times 10^{-35}$ |
| Vascular Progenitor cell:Blood | 94/100 | $1.75 \times 10^{-37}$ | $3.83 \times 10^{-35}$ |
| Neural Stem cell:Hippocampus | 92/99 | $1.33 \times 10^{-35}$ | $2.42 \times 10^{-33}$ |
| Neural Progenitor cell:Embryonic Prefrontal Cortex | 118/166 | $3.90 \times 10^{-23}$ | $6.08 \times 10^{-21}$ |
| Villous cytotrophoblast:Placenta | 72/99 | $1.61 \times 10^{-15}$ | $2.20 \times 10^{-13}$ |
| Lake Et al.Science.In8:Brain | 69/98 | $8.83 \times 10^{-14}$ | $1.07 \times 10^{-11}$ |
| T cell:Undefined | 67/95 | $1.77 \times 10^{-13}$ | $1.94 \times 10^{-11}$ |

Table S11: Top 10 enriched terms for the "Tabula Sapiens" category in Enrichr for HVG. Full list is available in the Supplementary Materials.

| Term | Overlap | $P$-value | Adjusted $P$-value |
|---|---|---|---|
| Large Intestine-monocyte | 60/100 | $5.11 \times 10^{-8}$ | $2.40 \times 10^{-5}$ |
| Small Intestine-monocyte | 55/100 | $8.01 \times 10^{-6}$ | $1.88 \times 10^{-3}$ |
| Trachea-neutrophil | 52/100 | $1.04 \times 10^{-4}$ | $1.63 \times 10^{-2}$ |
| Spleen-neutrophil | 48/100 | $1.87 \times 10^{-3}$ | $2.20 \times 10^{-1}$ |
| Bone Marrow-cd24 Neutrophil | 47/100 | $3.51 \times 10^{-3}$ | $2.35 \times 10^{-1}$ |
| Lung-neutrophil | 47/100 | $3.51 \times 10^{-3}$ | $2.35 \times 10^{-1}$ |
| Bone Marrow-nampt Neutrophil | 47/100 | $3.51 \times 10^{-3}$ | $2.35 \times 10^{-1}$ |
| Lymph Node-neutrophil | 45/99 | $8.94 \times 10^{-3}$ | $4.32 \times 10^{-1}$ |
| Lung-capillary Aerocyte | 45/100 | $1.11 \times 10^{-2}$ | $4.32 \times 10^{-1}$ |
| Muscle-capillary Endothelial Cell | 45/100 | $1.11 \times 10^{-2}$ | $4.32 \times 10^{-1}$ |

Table S12: Top 10 enriched terms for the "GO Biological Process 2026" category in Enrichr for TD and RNA-only. If they do not match, they are shown as [TD,RNA-only]. Full list is available in the Supplementary Materials.

| Term | Overlap | $P$-value | Adjusted $P$-value |
|---|---|---|---|
| RNA Processing (GO:0006396) | [234,207]/319 | $[1.54, 1.02] \times 10^{-[48,30]}$ | $[8.61, 1.15] \times 10^{-[45,27]}$ |
| Translation (GO:0006412) | [196,187]/251 | $[6.13, 1.33] \times 10^{-[48,40]}$ | $[1.71, 5.64] \times 10^{-[44,37]}$ |
| DNA Damage Response (GO:0006974) | [306,282]/464 | $[4.05, 6.49] \times 10^{-[47,34]}$ | $[7.54, 9.14] \times 10^{-[44,31]}$ |
| Protein Biosynthetic Process (GO:0160307) | [130,127]/147 | $[7.65, 2.00] \times 10^{-[44,40]}$ | $[1.07, 5.64] \times 10^{-[40,37]}$ |
| MRNA Processing (GO:0006397) | [205,165]/277 | $[1.21, 4.15] \times 10^{-[43,19]}$ | $[1.35, 1.37] \times 10^{-[40,16]}$ |
| Chromatin Remodeling (GO:0006338) | [201,185]/290 | $[5.95, 3.05] \times 10^{-[36,26]}$ | $[5.53, 1.91] \times 10^{-[33,23]}$ |
| Cytoplasmic Translation (GO:0002181) | 96/106 | $1.05 \times 10^{-34}$ | $[8.37, 1.91] \times 10^{-[32,31]}$ |
| DNA Repair (GO:0006281) | [216,198]/325 | $[2.75, 2.44] \times 10^{-[34,24]}$ | $[1.25]1.92 \times 10^{-[31,21]}$ |
| DNA Metabolic Process (GO:0006259) | [220,209]/336 | $[1.96, 2.53] \times 10^{-[33,27]}$ | $[1.21, 2.37] \times 10^{-[30,24]}$ |
| MRNA Splicing, via Spliceosome (GO:0000398) | [172,139]/242 | $[4.23, 1.59] \times 10^{-[33,14]}$ | $[2.36, 2.56] \times 10^{-[30,12]}$ |
| Macromolecule Biosynthetic Process (GO:0009059) | [165,163]/243 | $[4.12, 7.28] \times 10^{-[28,27]}$ | $[1.44, 5.85] \times 10^{-[25,24]}$ |
| Chromatin Organization (GO:0006325) | [231,220]/362 | $[1.71, 1.07] \times 10^{-[32,26]}$ | $[8.67, 7.56] \times 10^{-[30,24]}$ |
| Ribosome Biogenesis (GO:0042254) | [130,123]/171 | $[4.12, 8.29] \times 10^{-[27,25]}$ | $[1.64, 4.67] \times 10^{-[27,22]}$ |

Table S13: Top 10 enriched terms for the "GO Cellular Component 2026" category in Enrichr for TD and RNA-only. If they do not match, they are shown as [TD,RNA-only]. Full list is available in the Supplementary Materials.

| Term | Overlap | $P$-value | Adjusted $P$-value |
|---|---|---|---|
| Nucleus (GO:0005634) | [2407,2269]/5331 | $[2.45, 1.17] \times 10^{-[95,58]}$ | $[1.17, 5.57] \times 10^{-[92,56]}$ |
| Intracellular Membrane-Bounded Organelle (GO:0043231) | [2464,2330]/5509 | $[8.72, 7.12] \times 10^{-[93,58]}$ | $[2.08, 1.70] \times 10^{-[90,55]}$ |
| Nuclear Lumen (GO:0031981) | [449,414]/705 | $[3.04, 3.52] \times 10^{-[62,44]}$ | $[4.83, 4.20] \times 10^{-[60,42]}$ |
| Nucleolus (GO:0005730) | [433,400]/684 | $[6.59, 4.31] \times 10^{-[59,42]}$ | $[7.84, 4.12] \times 10^{-[57,40]}$ |
| Intracellular Membraneless Organelle (GO:0043232) | [662,637]/1196 | $[1.71, 1.62] \times 10^{-[57,47]}$ | $[1.62, 2.58] \times 10^{-[55,45]}$ |
| Nuclear Ribonucleoprotein Granule (GO:0140168) | [234,200]/364 | $[1.24, 2.26] \times 10^{-[33,17]}$ | $[9.84, 7.21] \times 10^{-[32,16]}$ |
| Nuclear Speck (GO:0016607) | [207,179]/320 | $[1.95, 1.02] \times 10^{-[30,16]}$ | $[1.32, 3.06] \times 10^{-[28,15]}$ |
| Focal Adhesion (GO:0005925) | [227,233]/371 | $[4.24, 3.14] \times 10^{-[28,31]}$ | $[2.52, 2.50] \times 10^{-[26,29]}$ |
| Cell-substrate Junction (GO:0030055) | [229,235]/379 | $[3.12, 2.81] \times 10^{-[27,30]}$ | $[1.65, 1/91] \times 10^{-[25,28]}$ |
| U2-type Spliceosomal Complex (GO:0005684) | [80,66]/93 | $[1.17, 1.67] \times 10^{-[25,13]}$ | $[5.57, 3.46] \times 10^{-[24,12}$ |
| Extracellular Exosome (GO:0070062) | [896,902]/2059 | $[1.55, 8.44] \times 10^{-[23,25]}$ | $[6.72, 5.04] \times 10^{-[22,23]}$ |
| Extracellular Vesicle (GO:1903561) | [898,904]/2082 | $[3.60, 2.16] \times 10^{-[22,23]}$ | $[1.43, 1.15] \times 10^{-[20,21]}$ |
| Large Ribosomal Subunit (GO:0015934) | 51/55 | $3.88 \times 10^{-20}$ | $[1.32, 1.86] \times 10^{-18}$ |

Table S14: Top 10 enriched terms for the "GO Molecular Function 2026" category in Enrichr for TD and RNA-only. If they do not match, they are shown as [TD,RNA-only]. Full list is available in the Supplementary Materials.

| Term | Overlap | $P$-value | Adjusted $P$-value |
|---|---|---|---|
| Ubiquitin-like Protein Ligase Binding (GO:0044389) | [183,154]/281 | $[1.39, 9.11] \times 10^{-[27,17]}$ | $[1.72, 2.89] \times 10^{-[24,14]}$ |
| Cadherin Binding (GO:0045296) | [201,202]/321 | $[7.32, 2.14] \times 10^{-27}$ | $[4.55, 2.71] \times 10^{-24}$ |
| MRNA Binding (GO:0003729) | [209,179]/339 | $[1.38, 1.50] \times 10^{-[26,13]}$ | $[5.71, 3.17] \times 10^{-[24,11]}$ |
| Ubiquitin Protein Ligase Binding (GO:0031625) | [170,154]/264 | $[6.81, 9.11] \times 10^{-[25,17]}$ | $[2.12, 2.89] \times 10^{-[22,14]}$ |
| Protein Serine/Threonine Kinase Activity (GO:0004674) | 209/381 | $5.50 \times 10^{-18}$ | $[1.37, 3.48] \times 10^{-15}$ |
| Ubiquitin-protein Transferase Activity (GO:0004842) | [216,187]/417 | $[6.82, 8.28] \times 10^{-[15,7]}$ | $[1.41, 5.19] \times 10^{-[12,5]}$ |
| Kinase Binding (GO:0019900) | [236,232]/467 | $[1.40, 2.45] \times 10^{-[14,13]}$ | $[2.49, 4.43] \times 10^{-[12,11]}$ |
| Nuclear Receptor Binding (GO:0016922) | [80,74]/118 | $[2.70, 7.93] \times 10^{-[14,11]}$ | $[4.20, 1.01] \times 10^{-[12,8]}$ |
| Single-stranded DNA Binding (GO:0003697) | [69,65]/100 | $[4.25, 1.18] \times 10^{-[13,10]}$ | $[5.87, 1.36] \times 10^{-[11,8]}$ |
| Protein Kinase Binding (GO:0019901) | [257,262]/531 | $[5.33, 1.91] \times 10^{-[13,14]}$ | $[6.62, 4.84] \times 10^{-[11,12]}$ |
| Adenyl Ribonucleotide Binding (GO:0032559) | 173/339 | $1.67 \times 10^{-11}$ | $[1.48, 2.64] \times 10^{-9}$ |
| ATP Binding (GO:0005524) | [158,156]/300 | $[5.21, 2.59] \times 10^{-[12,11]}$ | $[5.39, 3.64] \times 10^{-[10,9]}$ |
